# Advantages of genotype imputation with ethnically matched reference panel for rare variant association analyses

**DOI:** 10.1101/579201

**Authors:** Mart Kals, Tiit Nikopensius, Kristi Läll, Kalle Pärn, Timo Tõnis Sikka, Jaana Suvisaari, Veikko Salomaa, Samuli Ripatti, Aarno Palotie, Andres Metspalu, Tõnu Esko, Priit Palta, Reedik Mägi

## Abstract

Genotype imputation has become a standard procedure prior genome-wide association studies (GWASs). For common and low-frequency variants, genotype imputation can be performed sufficiently accurately with publicly available and ethnically heterogeneous reference datasets like 1000 Genomes Project (1000G) and Haplotype Reference Consortium panels. However, the imputation of rare variants has been shown to be significantly more accurate when ethnically matched reference panel is used. Even more, greater genetic similarity between reference panel and target samples facilitates the detection of rare (or even population-specific) causal variants. Notwithstanding, the genome-wide downstream consequences and differences of using ethnically mixed and matched reference panels have not been yet comprehensively explored.

We determined and quantified these differences by performing several comparative evaluations of the discovery-driven analysis scenarios. A variant-wise GWAS was performed on seven complex diseases and body mass index by using genome-wide genotype data of ∼37,000 Estonians imputed with ethnically mixed 1000G and ethnically matched imputation reference panels. Although several previously reported common (minor allele frequency; MAF > 5%) variant associations were replicated in both resulting imputed datasets, no major differences were observed among the genome-wide significant findings or in the fine-mapping effort. In the analysis of rare (MAF < 1%) coding variants, 46 significantly associated genes were identified in the ethnically matched imputed data as compared to four genes in the 1000G panel based imputed data. All resulting genes were consequently studied in the UK Biobank data.

These associations provide a solid example of how rare variants can be efficiently analysed to discover novel, potentially functional genetic variants in relevant phenotypes. Furthermore, our work serves as proof of a cost-efficient study design, demonstrating that the usage of ethnically matched imputation reference panels can enable substantially improved imputation of rare variants, facilitating novel high-confidence findings in rare variant GWAS scans.

**Author summary:** Over the last decade, genome-wide association studies (GWASs) have been widely used for detecting genetic biomarkers in a wide range of traits. Typically, GWASs are carried out using chip-based genotyping data, which are then combined with a more densely genotyped reference panel to infer untyped genetic variants in chip-typed individuals. The latter method is called genotype imputation and its accuracy depends on multiple factors. Publicly available and ethnically heterogeneous imputation reference panels (IRPs) such as 1000 Genomes Project (1000G) are sufficiently accurate for imputation of common and low-frequency variants, but custom ethnically matched IRPs outperform these in case of rare variants. In this work, we systematically compare downstream association analysis effects on eight complex traits in ∼37,000 Estonians imputed with ethnically mixed and ethnically matched IRPs. We do not observe major differences in the single variant analysis, where both imputed datasets replicate previously reported significant loci. But in the gene-based analysis of rare protein-coding variants we show that ethnically matched panel clearly outperforms 1000G panel based imputation, providing 10-fold increase in significant gene-trait associations. Our study demonstrates empirically that imputed data based on ethnically matched panel is very promising for rare variant analysis – it captures more population-specific variants and makes it possible to efficiently identify novel findings.

## Introduction

Genome-wide association studies (GWASs) have been successfully implemented to capture genetic variants with small to modest effect sizes and have identified thousands of common variants robustly associated with different complex traits and diseases [1]. However, even in aggregate, these explain only a small fraction of the heritability of studied diseases.

The sample size of a GWAS can be increased through relatively cheap chip-based genotyping and subsequent genotype imputation. Imputation is a commonly used computational method for lending information from a densely genotyped reference panel of phased haplotypes, allowing to study variants that have not been directly genotyped in target samples and thereby this approach not only increases the power but also the resolution of GWAS [2–4]. Genotype imputation can also facilitate better fine-mapping association signals through the increase of genetic variant density in candidate genomic regions [5].

Publicly accessible imputation reference panels like 1000 Genomes Project (1000G) [6] and Haplotype Reference Consortium (HRC) [7] have been frequently used for imputation in advance to GWAS. Nevertheless, both of these ethnically heterogeneous reference panels have only limited capacity to provide complete and accurate imputation of rare (minor allele frequency; MAF < 1%) variants [8], suggested to contribute to the missing heritability [9]. During the last few years it has been shown that using an ethnically matched reference panel can greatly improve the ‘completeness’ and accuracy of genotype imputation [10–15], resulting in higher imputation accuracy compared to the 1000G panel even in case of smaller panel size [16,17]. In addition, several recent studies have demonstrated the utility of ethnically matched datasets for the discovery of disease or trait-associated rare variants [18–25].

Imputed datasets based on ethnically matched reference panels are considered to be powerful tools to discover previously unidentified rare variants. However, the typical approaches for testing associations of genetic variants with phenotypes based on simple regression models, and are underpowered for rare variants in most studies due to their low frequencies and large numbers [26]. To overcome these issues, different methods have been proposed to increase statistical power in rare variant association studies, typically by combining information across multiple rare variants within a specific genomic region or functional unit (e.g. gene) [27–29]. Often these methods focus on certain categories of variation (e.g. missense or loss-of-function (LoF) variants) [30], and have been applied successfully in several studies [31–33]. Therefore, gene-based tests allow to capture the joint contribution of multiple rare variants, improve power and enable to identify novel disease associated genes encompassing putatively functional variants [34–37].

In the current study, we impute 51,886 chip-typed Estonians with both ethnically matched Estonian-Finnish (EstFin) and ethnically mixed 1000G imputation reference panels (IRPs) to determine and quantify the differences in analysis results of eight complex traits. In particular, we evaluate two analysis scenarios: 1) a variant-wise GWAS; 2) a gene-wise analysis to determine the joint contribution of rare (MAF < 1%) nonsynonymous (NS) and LoF variants which we validate in the UK Biobank data.

## Results

First, we developed a high-coverage (∼30×) whole genome sequencing (WGS) based imputation reference panel comprising of ethnically closely related 2,279 Estonians and 1,856 Finns, resulting in 8,270 haplotypes in the EstFin IRP. Secondly, we imputed 51,886 chip-genotyped Estonians with the EstFin and 1000G IRPs (S1 Appendix, S1 Table, S1 and S2 Figs). The EstFin IRP provided 13.86 million (M) and the 1000G IRP 9.06 M confidently imputed variants (imputation INFO-value > 0.8) with MAF > 0.05% in 36,716 unrelated individuals, which were further used to carry out a comparative GWAS and gene-wise association testing of rare variants with eight complex traits. Finally, the identified significant gene-trait associations were studied in the UK Biobank data (Fig 1).

**Fig 1.**
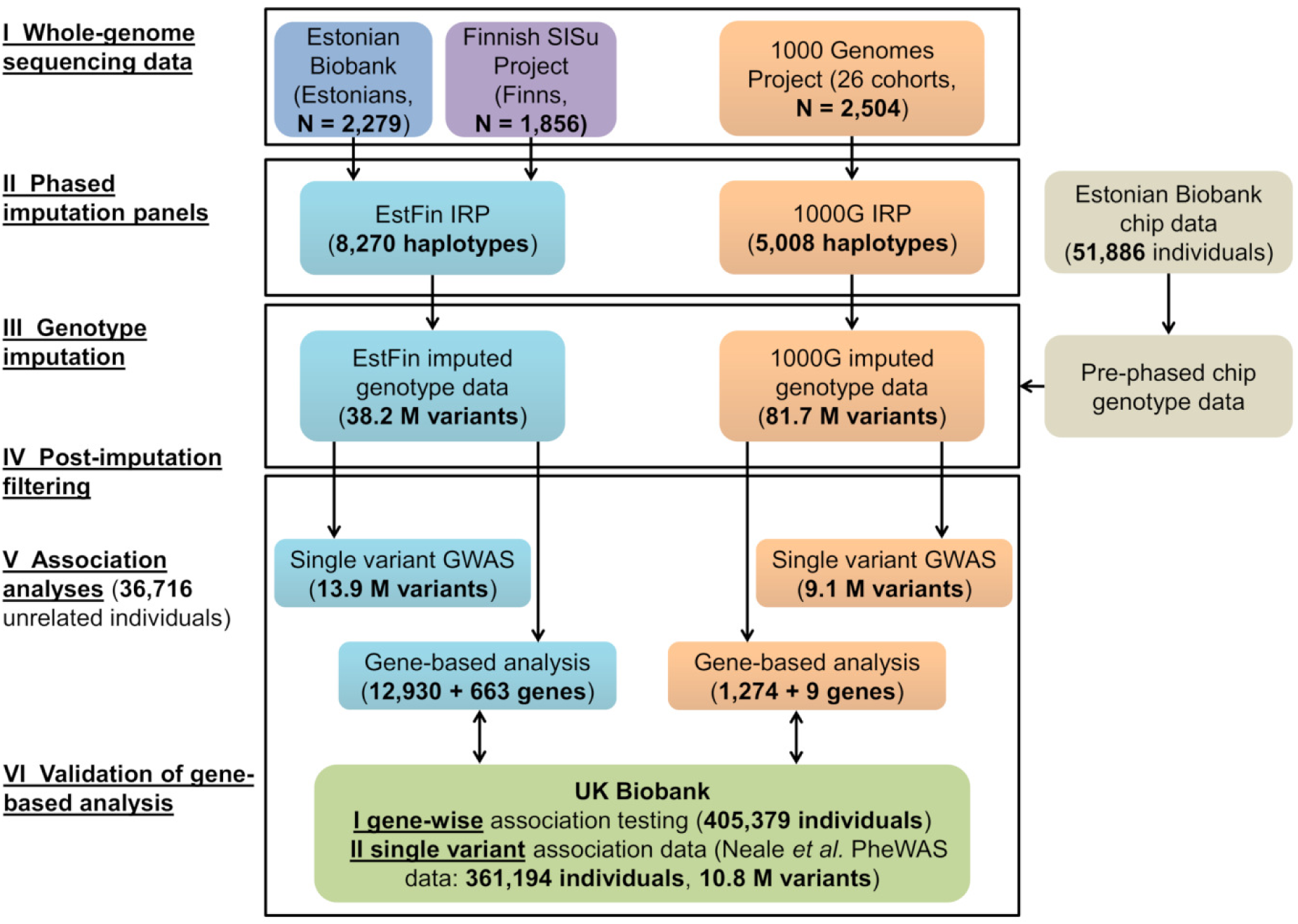
Schematic overview of used datasets and main processing and analysis steps. This scheme gives an overview of used imputation reference panels, chip-based and imputed genotype datasets, and comparative association analyses, for which gene-based results were validated in the UK Biobank data.

### Single variant analysis

Genome-wide association testing was performed separately in both imputed datasets with eight complex traits: body mass index and seven complex diseases of major public health importance [38] – bipolar disorder (BD), coronary artery disease (CAD), Crohn’s disease (CD), hypertension (HT), rheumatoid arthritis (RA), type 1 diabetes (T1D), and type 2 diabetes (T2D) (S2 Table). We detected 12 and 13 genome-wide significant (*P* < 6.25 × 10^−9^) loci based on EstFin and 1000G IRPs, respectively. Results of variant-wise GWA studies are summarized in Table 1 and S3 Fig. In both datasets we discovered eleven identical loci and three IRP-specific associations, all of which have been previously reported [1] (S3 Table). Autoimmune diseases RA and T1D demonstrated common variant associations in the HLA-region, BMI and T2D revealed long established association with *FTO* gene (Fig 2). Although lead variants did not overlap (except rs11102694 and rs9273363 with T1D), top hits from both datasets were in close proximity and in high linkage disequilibrium (S4 Table and S4 Fig).

**Table 1.**
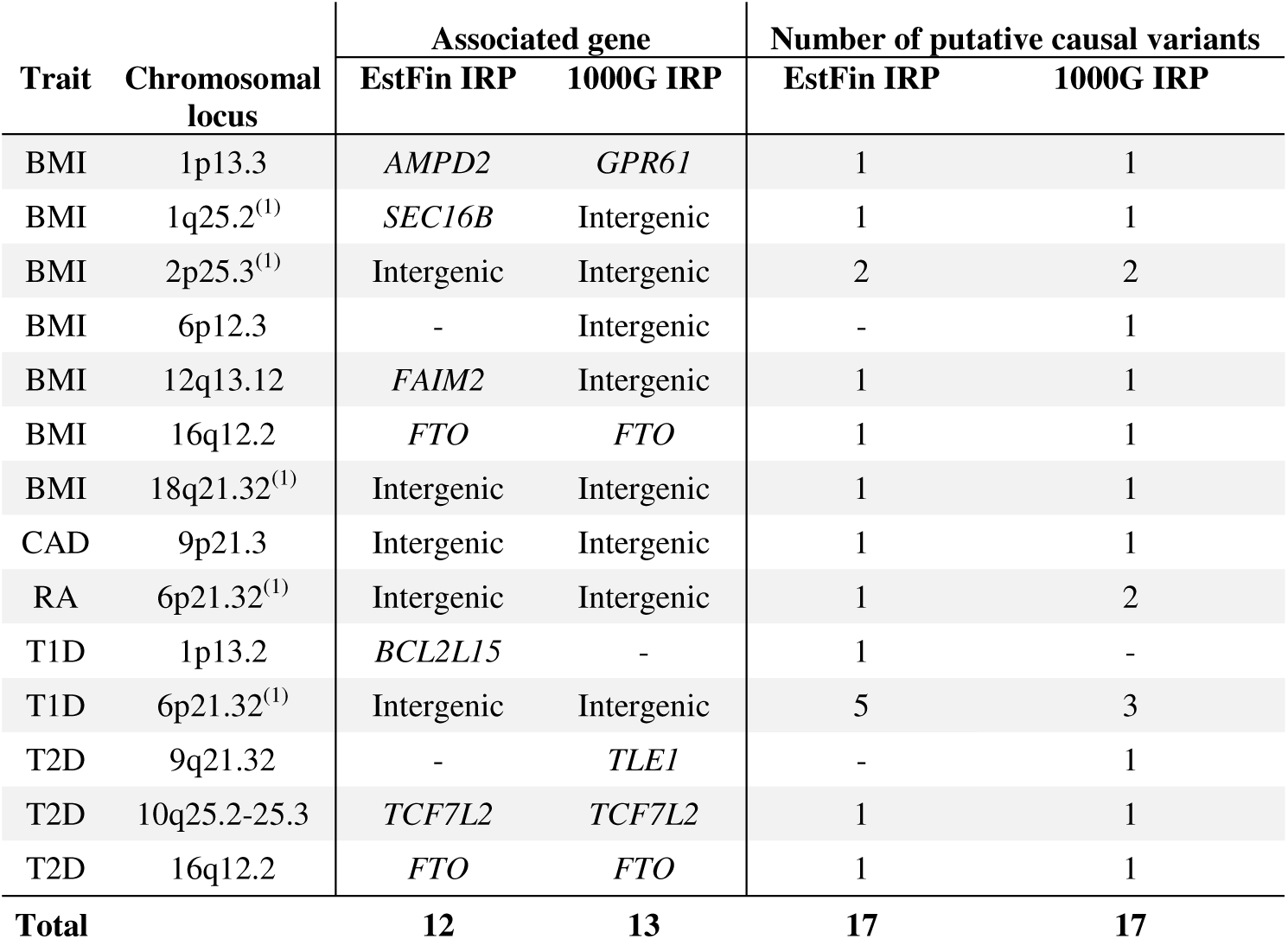
Study-wide significant loci detected in single variant association analysis. Confidently imputed variants (INFO > 0.8) were tested for associations with complex traits (BMI – body mass index, CAD – coronary artery disease, RA – rheumatoid arthritis, T1D – type 1 diabetes, T2D – type 2 diabetes). Analyses were conducted separately in the EstFin-based and the 1000G-based imputed datasets. The study-wide significance threshold after correction for multiple testing was set to *P* < 6.25 × 10^−9^ and for each locus, associated gene containing variant with the lowest *P* value was reported. In the last two columns, results of fine-mapping analysis are shown with the numbers of putative causal variants per genomic region. ^(1)^ Locus including significant (*P* < 3.27 × 10^−8^, 10-fold enrichment in Estonians) variants detected in MAF-enriched analysis in comparison of 503 European individuals from the 1000G phase 3 data.

**Fig 2.**
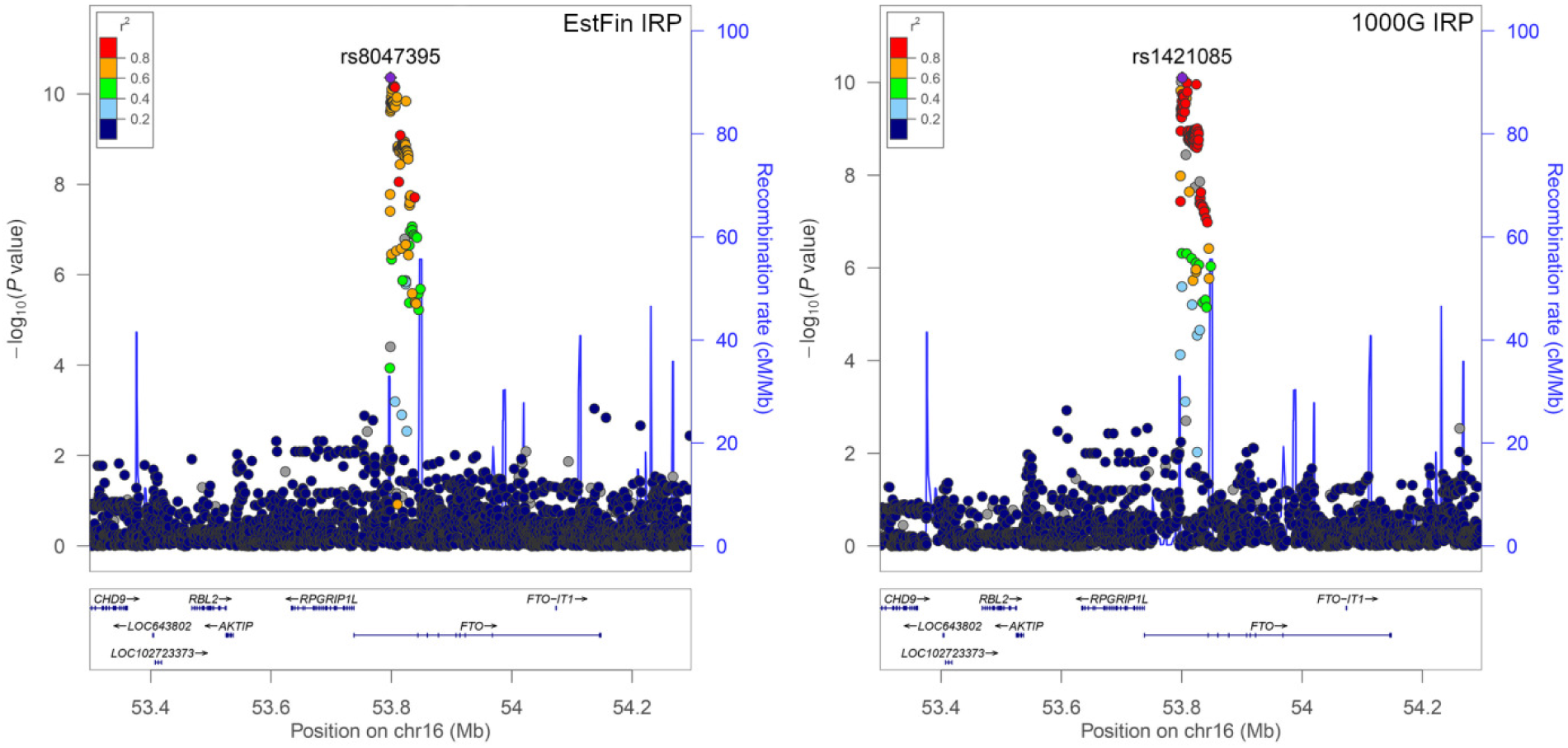
Regional association plots of type 2 diabetes at the *FTO* locus. The left panel of regional plot shows the genome-wide association analysis results for the EstFin-based imputed data, while the right panel shows results for the 1000G IRP imputed data. The purple symbol represents the lead variant, and the rest of the colour-coded variants denote LD with the lead variant estimated by *r*^*2*^ from the 1000G phase 3 (EUR population) data. Comparison of analysis results of both imputed datasets indicates that variants with the lowest *P* value do not overlap, but are highly correlated. Imputed data based on the EstFin panel provides evidence for an association between T2D and alleles of the *FTO* locus with the lowest *P* value at rs8047395 (*P* = 4.4 × 10^−11^). The same SNV shows a significant association in the data imputed with 1000G IRP (*P* = 1.5 × 10^−10^), but the lowest *P* value is at rs1421085 (P = 7.9 × 10^−11^, Pearson’s *r*^*2*^=0.82 between rs8047395 and rs1421085).

IRP-specific associations included *BCL2L15* association with T1D in case of the EstFin panel based imputation, whereas an intergenic locus at chromosome 6p12.3 was associated with BMI and *TLE1* locus with T2D in the 1000G-based imputed data only. Although, for both associations, the lowest *P* values in the other IRP-based imputed data were very close to the genome-wide significance level (S4 Table and S3 Fig).

To identify the likely causal variant at each locus, we performed fine-mapping analysis in all significant genomic regions discovered in genome-wide association scan. All but three (BMI at 2p25.3, RA and T1D at the HLA-region) significant regions demonstrated only one likely causal variant (Table 1). We also tested variants having allele frequency enrichment in Estonians as compared to the 503 European individuals from the 1000G data. Genome-wide significant (*P* < 3.27 × 10^−8^ for 10-fold enrichment) variants were detected for BMI at 1q25.2, 2p25.3, and 18q21.32 loci and for RA and T1D in the HLA-region in both imputed datasets (Table 1).

In conclusion, single variant association analyses did not indicate major differences in results based on data imputed with ethnically matched and mixed IRPs.

### Gene-based analysis

In addition to single variant analyses, we conducted gene-based association testing of rare (MAF < 1%) nonsynonymous (NS) and loss-of-function (LoF) variants for eight complex traits in both imputed datasets. When considering genes with at least two confidently imputed (INFO > 0.8) rare NS variants, we observed noteworthy differences in the number of genes analysed – EstFin IRP outperformed 1000G, providing 12,930 and 1,274 unique genes, respectively. In the analysis of rare LoF variants, we identified even more drastic differences – 663 genes were tested in the EstFin panel imputation and only six genes in the 1000G panel imputation.

Consequently, whilst testing the genes including rare NS variants, we detected 38 significant gene-trait associations (*P*_*NS*_ < 4.83 × 10^−7^) in the EstFin-based imputed data and four in the 1000G panel based imputation (Table 2). At significance level *P*_*LoF*_ < 9.40 × 10^−6^ we detected 10 genes including rare LoF variants based on the EstFin imputed data and none in the 1000G-based imputation (Table 2). Comparative results of gene-based analysis are presented in Figures 3 and S5. While none of these associated genes were implicated in our single variant GWAS, 122 NS and 22 LoF variants were involved in gene-wise analysis (S5 Table). We determined that the large majority (45 out of 52) of significant gene-trait associations relied on two or three NS/LoF variants. Seven out of 52 associations relied on four or more NS/LoF variants and the signal of joint contribution of rare variants was driven by multiple variants (*P* < 0.05) for 22/52 tests (S5 Table).

**Table 2.**
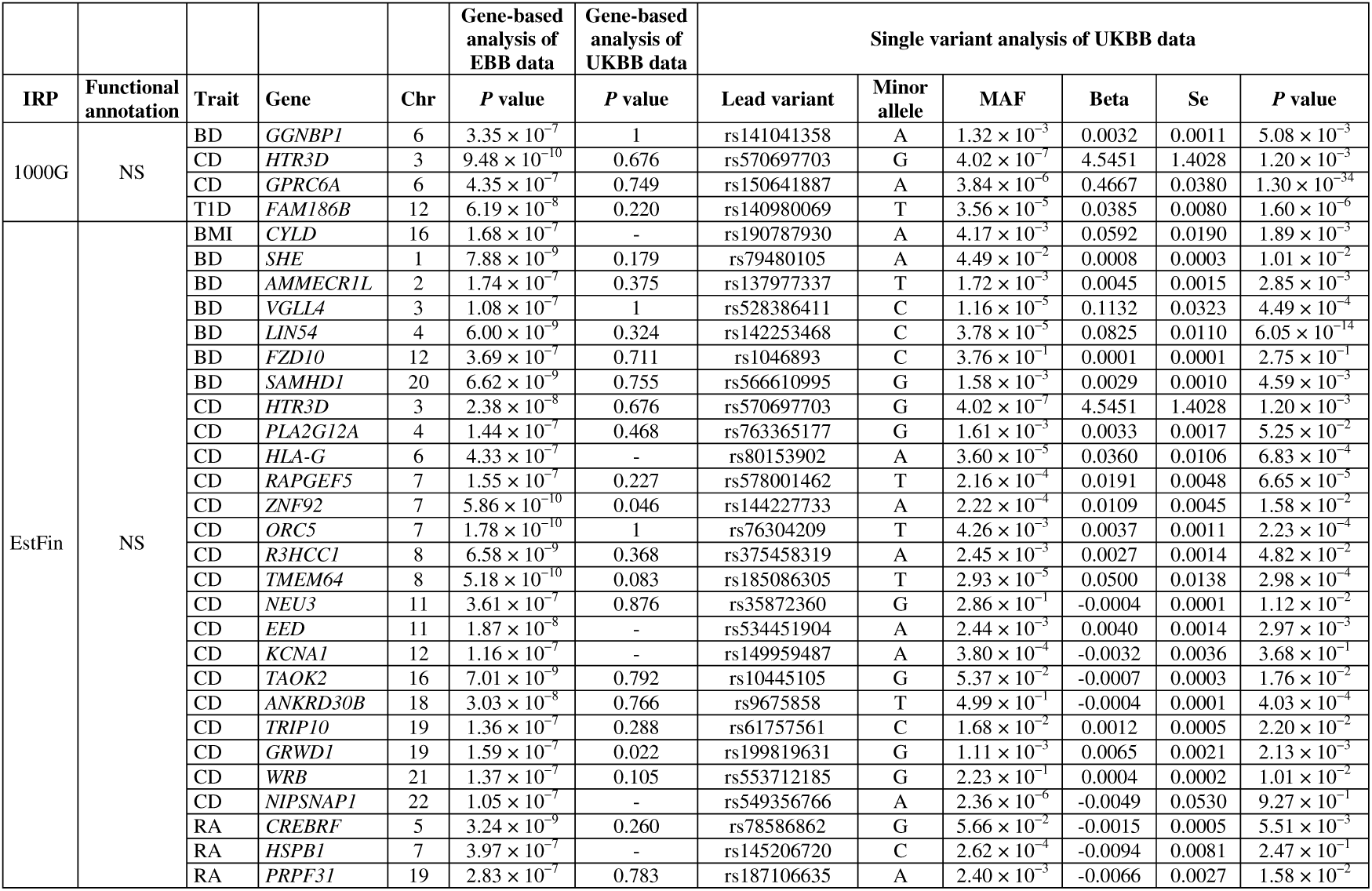
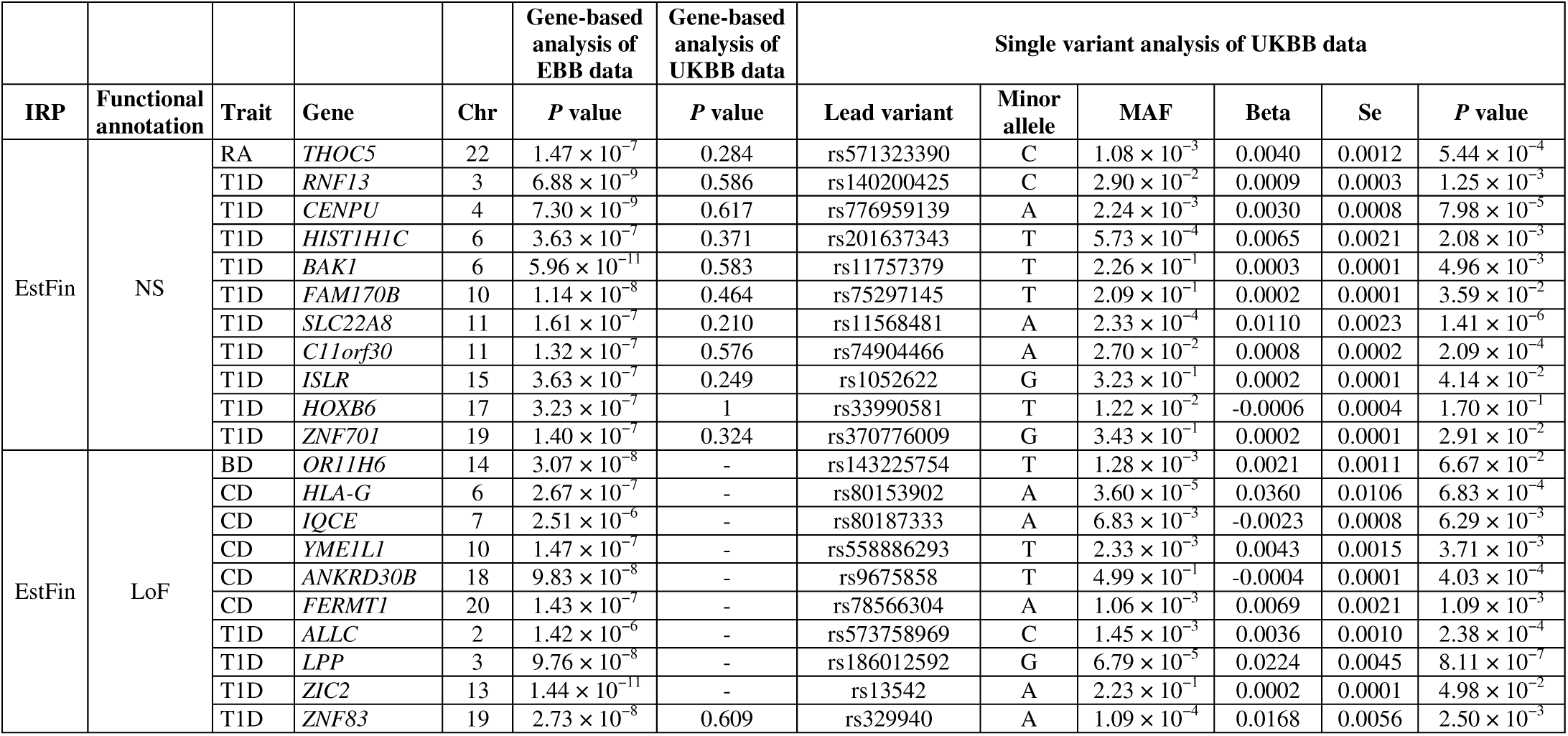
Identified significant gene-trait associations. Genes with at least two confidently imputed (INFO > 0.8) rare (MAF < 1%) nonsynonymous (NS) and loss-of-function (LoF) variants are tested for association with complex traits (BMI – body mass index, BD – bipolar disorder, RA – rheumatoid arthritis, T1D – type 1 diabetes, T2D – type 2 diabetes). Analyses are performed separately in the EstFin-based and the 1000G-based imputed datasets. Multiple testing corrected significance levels are applied based on the number of genes tested in both datasets: *P*_*NS*_ < 4.83 × 10^−7^ for genes containing NS variants and *P*_*LoF*_ < 9.40 × 10^−6^ for genes containing LoF variants. First, the results of gene-based analysis in 36,716 Estonian Biobank (EBB) individuals are presented. Next, the gene-trait associations are validated in 405,379 UK Biobank (UKBB) individuals. Finally, for each significant gene-trait result detected in the EBB data, variant-trait association with the smallest *P* value from the UKBB single variant GWAS is provided.

**Fig 3.**
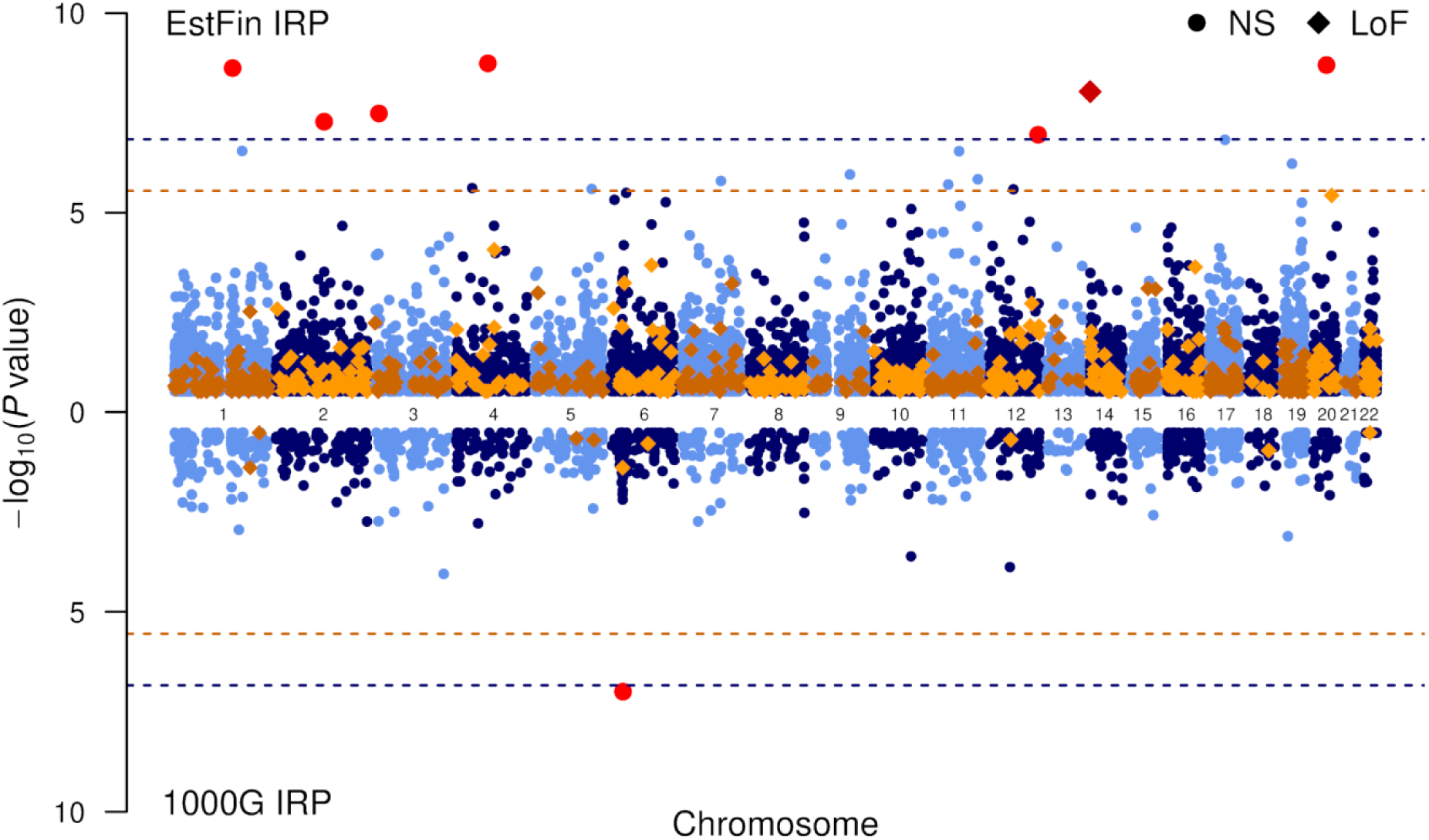
Miami plot of the gene-based association testing of bipolar disorder. The top panel shows the gene-based association analysis results using the EstFin-based imputed data, while the bottom part shows results for the 1000G IRP imputed data. Blue dots represent tested genes including NS variants and orange squares LoF variants. Dashed lines indicate significance levels after correction for multiple testing: *P* < 4.83 × 10^−7^ for NS variants (blue) and *P* < 9.40 × 10^−6^ for LoF variants (orange). Red symbols denote significant genes. In the analysis of NS variants we identify six genes based on the EstFin IRP data and one significant gene in the 1000G-based data. In the EstFin-based imputed data we detect a single gene-trait association of LoF substitutions, whereas none of the significant associations is observed in the data based on ethnically mixed 1000G IRP.

### Validation of gene-based analysis

First, all 52 unique genes that demonstrated significant gene-trait associations in gene-wise analysis were selected and gene-based association testing was repeated using 405,379 individuals from the UK Biobank [39,40] (Table 2). The strongest gene-trait associations were detected with CD in *GRWD1* (*P* = 0.022) and *ZNF92* (*P* = 0.046) genes, but neither of these were significant after correcting for multiple testing.

Secondly, we used variant-wise GWAS results of the UK Biobank in 361,194 individuals [41]. Although approximately only one fourth of the tested rare NS and LoF variants overlapped between the Estonian Biobank and the UK Biobank data, the UKBB GWAS analysis confirmed association signals (P < 10^−5^) in five detected genes. Considering the lowest *P* values in our candidate gene regions, we detected two significant (*P* = 1.30 × 10^−34^, CD in *GPRC6A* with the 1000G imputation; *P* = 6.05 × 10^−14^, BD in *LIN54* with the EstFin imputation) and three suggestive (*P* = 8.11 × 10^−7^, T1D in *LPP* with the EstFin imputation, *P* = 1.41 × 10^−6^, T1D in *SLC22A8* with the EstFin imputation, *P* = 1.60 × 10^−6^, T1D in *FAM186B* with the 1000G imputation) associations in the UKBB GWAS data (Table 2). Significant gene-based findings with plausible evidence of biological meaningfulness including *VGLL4, LPP* and *HLA-G* as well as few other loci are discussed in S1 Appendix. Relevant GWAS Catalog entries related to significant findings from gene-based analysis are presented in S6 Table.

Gene-based analysis of rare variants demonstrated that our ethnically matched panel clearly outperformed the 1000G-based imputation, providing approximately 10-fold increase in the number of tested genes and significant findings. Further literature and GWAS Catalog mining of these significant associations determined that most of these associated genes have been previously indicated, but there were also several novel findings.

## Discussion

Over the past few years, several ethnically matched imputation reference panels have been developed to complement the widely used cosmopolitan 1000G and HRC panels. The former panels have showed great improvement in imputation accuracy, but their effect to the downstream analyses have not been well examined. In the current study, we performed a comparison of ethnically matched EstFin and ethnically mixed 1000G-based imputed genotypes in the Estonian Biobank study cohort of ∼52,000 individuals. In addition to the single variant analysis, we examined downstream differences as a measure of identified associations in MAF-enriched and rare-variant analysis. We demonstrate that ethnically matched reference panel empowers the detection of rare variant signals, enabling to identify clinically significant novel loci for complex diseases that will be further discussed below.

### Ethnically matched reference panel lead to considerable gain in downstream analyses for rare variants

Ethnically matched panel provides a significantly higher proportion of confidently imputed variants compared to the 1000G panel (S1 Table). The difference increases with the decrease of MAF, because of the insufficient representation of rare (or even population-specific) variation in the mixed-ethnicity panels like 1000G IRP. Correspondingly, we observed that single variant GWAS identified a similar number of genome-wide significant findings in these two imputed datasets and we did not detect any major differences in fine-mapping of these loci. At the same time, it should be taken into consideration that in the current analyses we rely on a relatively small number of disease cases, resulting in limited statistical power. It is likely one of the main factors why we did not observe any major differences in single variant GWA analyses. Possibly these results would be different with significantly larger cohorts as, at some point, one should start detecting low-frequency and rare variants that have been imputed confidently and therefore can be tested with the ethnically matched IRPs. Secondly, 1000G reference panel contains European haplotypes, and therefore it can be a relatively good imputation reference for common variation present in the Estonian population. But the results can differ for those populations, which are more distant from the populations used in transethnic imputation reference sets.

Rare variant analyses demonstrated great differences, where the EstFin-based imputation clearly out-performed the 1000G imputation, allowing for the identification of 10 times more genome-wide significant genes (Table 2). The EstFin IRP includes a larger number of haplotypes close to the target samples, deriving unique variants from genomes not included in the 1000G panel. This improves the chances of a rare variant being effectively tagged by a haplotype. Moreover, including haplotypes from ethnically distant populations may not accurately capture LD patterns of population-specific variants or imputation can introduce polymorphic variants in the target samples that are actually monomorphic as observed previously [10]. Our results empirically demonstrate the contribution of rare variants in complex traits analysis using ethnically matched panel, as compared to ethnically mixed population reference.

### Validation of gene-based analysis

Validation of gene-based analysis results in the UK Biobank individual-level data detected two significant gene-trait signals (P < 0.05), but neither remained significant after multiple testing correction. For gene-trait associations detected in the EBB data, we were not able to validate 6 (out of 42) and 9 (out of 10) genes containing NS and LoF variants, respectively. This was accounted for the UKBB data containing less than two NS and LoF variants within these genes (Table 2). A likely explanation is that the ethnically matched panel captures a significantly larger number of such rare variants which are not well-captured through the imputation with more heterogeneous reference panels. Therefore we argue that failing to validate most of the gene-based analysis results in the UK Biobank data can be due to the population-specific nature of the rare variant findings.

Nevertheless, some of the associations were validated by matching the observed significant genes with the variants located in the same gene regions in the UKBB single variant association analysis, as well as many of the genes detected by us were associated with relevant traits in literature (S1 Appendix). We hypothesize about a causative role for *LPP* variants conferring susceptibility to T1D – an assumption being initially rejected in a study involving both celiac disease and T1D patients [42]. Pleiotropic effects have been reported for *LPP* in association studies involving diverse autoimmune diseases where shared susceptibility factors outside the HLA-region are widely recognized. In addition, *LPP* mRNA and protein are expressed in multiple tissues, including islets of Langerhans and pancreas, and *LPP* gene is relatively intolerant of LoF variation (ExAC pLI = 0.58) [43].

In conclusion, ethnically matched IRPs enable considerably more confident imputation of low-frequency and rare variants as compared to ethnically mixed IRPs. Due to the size of our study dataset (36,716 individuals), small number of disease cases and consequently limited statistical power, we did not observe major differences in single variant GWAS results. However, in the gene-based analysis we observed a substantial advantage of ethnically matched IRP-based imputation over unmatched mixed reference panel-based imputation, enabling significantly improved gene-based analysis of also low-frequency and rare variants, allowing more efficient discovery of rare disease-associated genes and variants.

## Materials and methods

### Study cohorts

#### Estonian Biobank

The Estonian Biobank (EBB) is a population-based biobank of the Estonian Genome Center at the University of Tartu. EBB contains almost 52,000 individuals of the Estonian population (aged ≥18 years), which closely reflects the age, sex and geographical distribution of the Estonian adult population [44]. At baseline, the general practitioners performed a standardized health examination of the participants, who also donated blood samples for DNA, white blood cells and plasma tests and filled out a questionnaire on health-related topics. All biobank participants have signed a broad informed consent form, which allows periodical linking to national registries, electronic health record databases and hospital information systems. The majority of biobank participants have been analysed using genotyping arrays. High-coverage whole genome sequencing data is available for the 2,535 individuals, selected randomly by county of birth. The project was approved by the Research Ethics Committee of the University of Tartu (application number 234/T-12).

#### FINRISK

FINRISK is a series of health examination surveys carried out by the National Institute for Health and Welfare (formerly National Public Health Institute) of Finland every five years since 1972. The surveys are based on random population samples from five (or six in 2002) specified geographical areas of Finland. The samples have been stratified by 10-year age group, sex and study area. The sample sizes have varied from approximately 7,000 to 13,000 individuals and the participation rates from 60% to 90% in different study years. The age-range was 25-64 years until 1992 and 25-74 since 1997. The survey included a self-administered questionnaire, a standardized clinical examination carried out by specifically trained study nurses and drawing of a blood sample. Details of the examination have been previously described [45,46]. DNA has been collected since the 1992 survey from approximately 34,000 participants. The surveys have appropriate ethical approvals following the usual practices of each survey-year and the participants have signed an informed consent. The validity of clinical diagnoses in these registers has been documented in several publications [47–50].

#### Finnish Migraine Families collection

The families were collected over a period of 25 years from six headache clinics in Finland (Helsinki, Turku, Jyväskylä, Tampere, Kemi, and Kuopio) and through advertisements on the national migraine patient organization web page (http://migreeni.org/). Geographically, family members are represented from across the entire country. The current collection consists of 1,589 families, which included a complete range of pedigree sizes from small to large (e.g., 1,023 families had 1–4 related individuals and 566 families had 5+ related individuals). Currently, the collection consists of 8,319 family members, of whom 5,317 have a migraine diagnosis based on the third edition of the established International Classification for Headache Disorders (ICHD-3) criteria [51].

#### MESTA

The Living Conditions and Physical Health of Outpatients with Schizophrenia study recruited 276 outpatients with schizophrenia spectrum disorder (ICD-10 F20–F29) from the psychosis outpatient clinics of three municipalities in Finland (Järvenpää, Mäntsälä, and Tuusula). The study protocol consisted of a questionnaire and interview assessing current symptoms, functioning, lifestyle, and a comprehensive health examination. DNA samples were collected as a part of the study based on a separate informed consent [52,53].

#### Health 2000

The Finnish Health 2000 Survey was based on a nationally representative sample of 8,028 persons aged 30 years or over living in mainland Finland. A two-stage stratified cluster sampling design was used. The sampling frame was regionally stratified according to the five university hospital regions, and from each university hospital region 16 health care districts were sampled as clusters (altogether 80 health care districts). Persons within the health care districts were selected by systematic sampling, and persons aged 80 years and over were oversampled by doubling the sampling fraction. The field work took place between September 2000 and June 2001, and consisted of a home interview and a health examination at the local health centre, or a condensed interview and health examination of non-respondents at home. In addition, several questionnaires were used to assess symptoms, lifestyle, and exposures related to different health problems. Of the study sample, 88% were interviewed, 80% attended a comprehensive health examination and 5% attended a condensed examination at home [54].

### Ethnically matched imputation reference panel

WGS data for Estonian and Finnish samples were generated and jointly processed at the Broad Institute of MIT and Harvard. WGS samples had PCR-free DNA preparation (Estonian Biobank, FINRISK, Finnish Migraine Families collection, Health 2000) and PCR-amplified preparation (MESTA), followed by sequencing on the Illumina HiSeq X platform with the use of 151-bp paired-end reads with mean coverage of ∼30×. Sequenced reads were aligned to the GRCh37 human reference genome assembly using BWA-MEM v0.7.7 [55]; PCR duplicates were marked using Picard v1.136 (http://broadinstitute.github.io/picard/), and the Genome Analysis Toolkit (GATK) v3.4-46 [56,57] best-practice guideline was applied for further BAM processing and variant calling.

Samples were excluded based on high contamination (>5%), high proportion of chimeric alignment (>5%), low genotype quality (GQ < 50), low coverage (<20×), high coverage (> mean + 3 sd), relatedness (identity-by-descent (IBD) > 0.1), sex mismatches, high genotype discordance (>5%) between sequenced and chip-based data. Additionally, samples were filtered (mean ± 3 sd) based on total number of variants, non-reference variants, singletons, heterozygous/homozygous variants ratio (single nucleotide variants (SNVs) and indels were tested separately in the above-mentioned cases), insertion/deletion ratio for novel indels, insertion/deletion ratio for indels observed in dbSNP, and transition/transversion ratio. After filtering and exclusion of duplicates, the WGS datasets were merged, containing 4,135 individuals (2,279 Estonians and 1,856 Finns).

The following variants were set to missing: GQ < 20, read depth > 200×, phred-scaled genotype likelihood of reference allele < 20 for heterozygous and homozygous variant calls, and allele balance <0.2 or >0.8 for heterozygous calls. The GATK Variant Quality Score Recalibration (VQSR) was used to filter variants with a truth sensitivity of 99.8% for SNVs and of 99.9% for indels. Variants with inbreeding coefficient < –0.3, quality by depth < 2 for SNVs and < 3 for indels, call rate < 90%, and Hardy-Weinberg equilibrium (HWE) *P* value < 1×10^−9^ were removed. Monomorphic, multi-allelic variants, and low-complexity regions [58] were further excluded. The final IRP contains 38,226,084 variants.

Autosomal chromosomes and GRCh37 (hg19) human reference genome assembly was used for all analysis.

### Chip-based genotype data

The EBB participants have been analysed using Illumina genotyping arrays: 1) Global Screening Array (GSA, N=33,277), 2) HumanCoreExome (CE, N=7,832), 3) HumanOmniExpress (OMNI, N=8,137), and 4) 370K (N=2,640). Individuals with missing phenotype data were excluded. Final set of genotyped data contained 48,163 unique individuals. The genotype calling for the microarrays was performed using Illumina’s GenomeStudio v2010.3 software. The genotype calls for rare variants on the GSA array were corrected using the zCall software (version May 8th, 2012). After variant calling, the data was filtered using PLINK v.1.90 [59] by sample (call rate > 95%, no sex mismatches between phenotype and genotype data, heterozygosity < mean ± 3 sd) and marker-wise (HWE *P* value > 1 × 10^−6^, call rate > 95%, and for the GSA array additionally by Illumina GenomeStudio GenTrain score > 0.6, Cluster Separation Score > 0.4). Before the imputation, variants with MAF < 1% and C/G or T/A polymorphisms as well as indels were removed, as these genotype calls do not allow precise phasing and imputation.

### Phasing and imputation

The WGS-based imputation reference panel was phased using Eagle v2.3 [60,61] with default parameters except the *Kpbwt* parameter that was set to 20000 to increase accuracy. Pre-phasing of genotyped data was performed in similar manner for all four arrays separately with Eagle and imputed with Beagle v4.1 [62]. All pre-phased genotype datasets were imputed twice using the following reference panels: 1) EstFin IRP containing 8,270 reference haplotypes and 38.2 M autosomal variants; 2) 1000G IRP holding 5,008 reference haplotypes and 81.7 M autosomal markers. All four imputed arrays were merged by IRP with BCFtools v1.6 (https://samtools.github.io/bcftools/bcftools.html). Imputation information measure (INFO-value) [4] were added using BCFtools plugin ‘impute-info’. Monomorphic, multi-allelic and directly genotyped variants were excluded for all downstream analyses. Only confidently imputed variants (INFO-value > 0.8) with MAF > 0.05% were considered: 13,859,717 (12,872,515 SNVs, 987,202 indels) variants imputed with the EstFin IRP and 9,058,236 (8,232,261 SNVs, 825,975 indels) variants imputed with the 1000G IRP (Fig 1).

### Phenotypes

In the association analysis, only unrelated individuals were included (IBD sharing < 0.2). Samples were excluded by choosing the minimal list of related individuals to break all kinship ties and, if possible, cases were preferred over controls using RELOUT5 tool from Allele (http://www.toomashaller.com/allele.html). Questionnaire-based data was linked to the electronic health records (the Estonian Health Insurance database, data available for years 2003–2015) and other health-related databases like the Estonian Causes of Death Registry (2003–2015), and the Estonian Cancer Registry (2003–2013).

After linking, dead people without time of death, participants without records from registries, and individuals older than 80 years at recruitment were excluded. The latter because diagnoses in the elderly people are often related to significant risk-altering comorbidities (cancer or cardiovascular diseases). Associations of body mass index, three cardiometabolic (coronary artery disease, hypertension, type 2 diabetes) and four autoimmune (bipolar disorder, Crohn’s disease, rheumatoid arthritis, type 1 diabetes) diseases were analyzed in 36,716 Estonians (S2 Table).

### Single variants analysis

Single variant analysis was conducted with Hail 0.1 (http://broadinstitute.github.io/picard/). Linear regression was used to test each variant’s allelic dosage additive effect with body mass index, and Firth [63] logistic regression with seven diseases. Models were adjusted for age, sex, first ten principal components (PC1-10), and genotype array. Only confidently imputed variants (INFO > 0.8) with MAF > 0.05% were considered. A multiple testing corrected significance level (5 × 10^−8^ / 8 phenotypes) = 6.25 × 10^−9^ were used.

All genome-wide significant loci were visualized by regional association plots using LocusZoom v0.4.8 [64] with the 1000G phase 3 European population LD reference panel. Pairwise examination of quantile-quantile plots of GWAS *P* values indicated that the distribution of the test statistics were nearly identical for both datasets, and did not demonstrate significant genomic inflation (S6 Fig). All significantly associated loci were compared to the National Human Genome Research Institute (NHGRI-EBI) GWAS Catalog [1] (April 10, 2018) data.

### Fine-mapping

To identify causal variants that denote molecular mechanisms behind the associations, we performed fine-mapping analysis using FINEMAP v1.3 [65] around (± 500 kilobase (kb)) genome-wide significant loci detected by variant-wise analysis (Table 1). FINEMAP was applied with default parameters, allowing for at most five causal variants and the highest posterior probability for the number of causal signals was used.

### Enrichment analysis

Enriched variants in the EstFin imputed data were detected in comparison of 503 European (EUR) individuals from the 1000G phase 3 data. Enrichment rates were calculated as MAF in Estonians (Est) divided by MAF in 1000G EUR individuals:

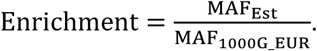

Corresponding Bonferroni corrected significance level for variants enriched 10-fold in Estonians (enrichment > 10) was [0.05 / (191,099 variants × 8 phenotypes)] = 3.27 × 10^−8^.

### Gene-based analysis

To determine the joint contribution of rare variants on eight complex traits, we implemented gene-based SKAT-O [27] tests with EPACTS (https://genome.sph.umich.edu/wiki/EPACTS). All variants were annotated with EPACTS module ‘anno’ (GENCODE v14 [66]). Only nonsynonymous (nonsynonymous, normal splice site or stop gain) and loss-of-function (stop gain, essential splice site or frameshift) variants with INFO > 0.8 and 0.000001% < MAF < 1% were included. Models were adjusted for age, sex, PC1-10 and genotype arrays. Results were post-filtered that each gene contained at least two NS or LoF variants. We identified 12,930 (663) and 1,274 (9) NS (LoF) genes in the EstFin and the 1000G IRP-based imputed data, respectively. Bonferroni corrected significance levels based on the number of identified genes in both imputed datasets were used: [0.05 / (12,951 × 8 phenotypes)] = 4.83 × 10^−7^ and [0.05 / (665 × 8 phenotypes)] = 9.40 × 10^−6^ for NS and LoF genes, respectively.

### UK Biobank data

The UK Biobank enrolled about 500,000 people aged between 40-69 years in 2006-2010 from across the United Kingdom [67]. Two approaches were used to validate significant gene-trait associations from the UKBB data:

1. We used genotyped and imputed individual-level data as released by UK Biobank in March 2018 [39,40]. In the analysis we used 405,379 unrelated (IBD sharing < 0.2) individuals with European origin and confidently imputed variants (INFO > 0.8). Diagnosis of prevalent disease was based on International Classification of Diseases (ICD-10) diagnosis codes and self-reported data. The same SKAT-O models were applied as used to discover 52 significant gene-trait associations in the Estonian Biobank data (Table 2).
2. We used GWAS analysis results of the UK Biobank in 361,194 individuals provided by the Neale lab [41] and selected variant-trait results with the lowest *P* value for each significant gene-trait association (gene ± 5 kb) detected in the Estonian Biobank data (Table 2).

## Acknowledgments

This research is financially supported by EU H2020 grant 692145, the European Regional Development Fund, Center of Excellence in Genomics GENTRANSMED (Project No. 2014-2020.4.01.15-0012), the Estonian Research Council Grants PUTJD817, IUT20-60, OUT1665P. We would like to acknowledge the International SISu Project for sharing the Finnish anonymised imputation reference panel data. We would like to acknowledge the High Performance Computing Center of the University of Tartu. PP and KP were supported by the NIASC - Nordic Information for Action eScience Center (a Nordic Center of Excellence; financed by NordForsk; Project No. 62721) grant to AP and SR. We would like to thank William Rayner for providing necessary files for genotype data preparation in his website. We would like to thank the UK Biobank for sharing the data (application No. 17085).

## Supporting information

**S1 Fig. Number of confidently imputed variants.** Venn diagrams of confidently imputed variants (INFO > 0.8) using EstFin (blue) and 1000G (orange) IRPs in four minor allele frequency categories. **(A)** 0.05% < MAF ≤ 0.5%, **(B)** 0.5% < MAF ≤ 1%, **(C)** 1% < MAF ≤ 5%, **(D)** MAF > 5%.

**S2 Fig. Distributions of imputation INFO-values.** Distribution of imputation INFO-values for the EstFin and the 1000G IRPs imputed data measured on chromosome 20 in four minor allele frequency categories. **(A)** EstFin IRP, **(B)** 1000G IRP.

**S3 Fig. Single variant association analysis results.** The top panel of Miami plot shows the single variant association analysis results using the EstFin-based imputed data, while the bottom part shows GWAS results for the 1000G IRP imputed data. Red dots denote significant regions and the genome-wide significance threshold after correction for multiple testing (*P* < 6.25 × 10^−9^) is indicated by a red dashed line.

**S4 Fig. Significant loci from single variant association analysis.** Regional plots show all significant genomic regions from the single variant analysis. The left panel indicates results for the EstFin-based imputed data, while the right panel shows results for the 1000G IRP imputed data. The purple symbol represents the lead variant, and the rest of the colour-coded variants denote LD with the lead variant estimated by *r*^*2*^ from the 1000G phase 3 (EUR population) data.

**S5 Fig. Gene-based association analysis results.** The top panel of Miami plot shows the gene-based association analysis results using the EstFin-based imputed data, while the bottom part shows results for the 1000G IRP imputed data. Blue dots represent tested genes including NS variants and orange asterisks LoF variants. Dashed lines indicate significance levels after correction for multiple testing: *P* < 4.83 × 10^−7^ for NS variants (blue) and *P* < 9.40 × 10^−6^ for LoF variants (orange). Red symbols denote significant genes.

**S6 Fig. Quantile-quantile plots for single variant association analysis.** Quantile-quantile plots of the GWAS *P* values based on the EstFin (left panel, blue dots) and the 1000G (right panel, purple dots) IRPs imputed data. Region in gray dashed lines is the 95% confidence band.

**S1 Table. Number of imputed variants.** Number of overall, well-imputed (INFO > 0.4) and confidently imputed (INFO > 0.8) variants with the EstFin and the 1000G IRPs. The last column indicates confidently imputed variants common for both IRP-based imputations.

**S2 Table. An overview of seven complex diseases.** ICD-10 diagnosis codes, number of cases and controls for seven complex diseases in 36,716 Estonian Biobank individuals used in the association analysis.

**S3 Table. Previously known associations for significant single variant analysis results.** Relevant genome-wide associations from the GWAS Catalog for significant genomic regions (around genes (± 50 kb) with the lowest *P* value) detected by variant-wise analysis. Gray background refers to direct relationship between studied trait and GWAS Catalog entry.

**S4 Table. Summary statistics of lead variants at significant loci identified in single variant GWAS results.** Confidently imputed variants (INFO > 0.8) are tested for associations with complex traits (BMI – body mass index, CAD – coronary artery disease, RA – rheumatoid arthritis, T1D – type 1 diabetes, T2D – type 2 diabetes). Analyses are conducted separately for the EstFin and the 1000G IRP-based imputed datasets. Multiple testing corrected significance level (*P* < 6.25 × 10^−9^) is used. For each significant loci, lead variant with single variant GWAS summary statistics are provided.

**S5 Table. Overview of genetic variants involved in significant gene-trait associations.** A list of all single variants involved in significant gene-wise associations with single variant GWAS summary statistics. In the last column, relevant references are provided, where particular gene-trait association is previously identified.

**S6 Table. Previously known associations for significant gene-based analysis results.** Relevant genome-wide associations from the GWAS Catalog for significant genes (± 50 kb) detected by gene-wise analysis. Gray background refers to direct relationship between studied trait and GWAS Catalog entry.

**S1 Appendix. Overview of genotype imputation and examples of disease associated genes.** A detailed overview of genotype imputation with the EstFin and the 1000G IRPs are provided. Five examples of significant gene-trait associations from the EstFin-based imputation results with potential underlying biological mechanisms are considered in more details.

