## Supplementary figures and images for "Advantages of genotype imputation with ethnically matched reference panel for rare variant association analyses"

### S1 Figure

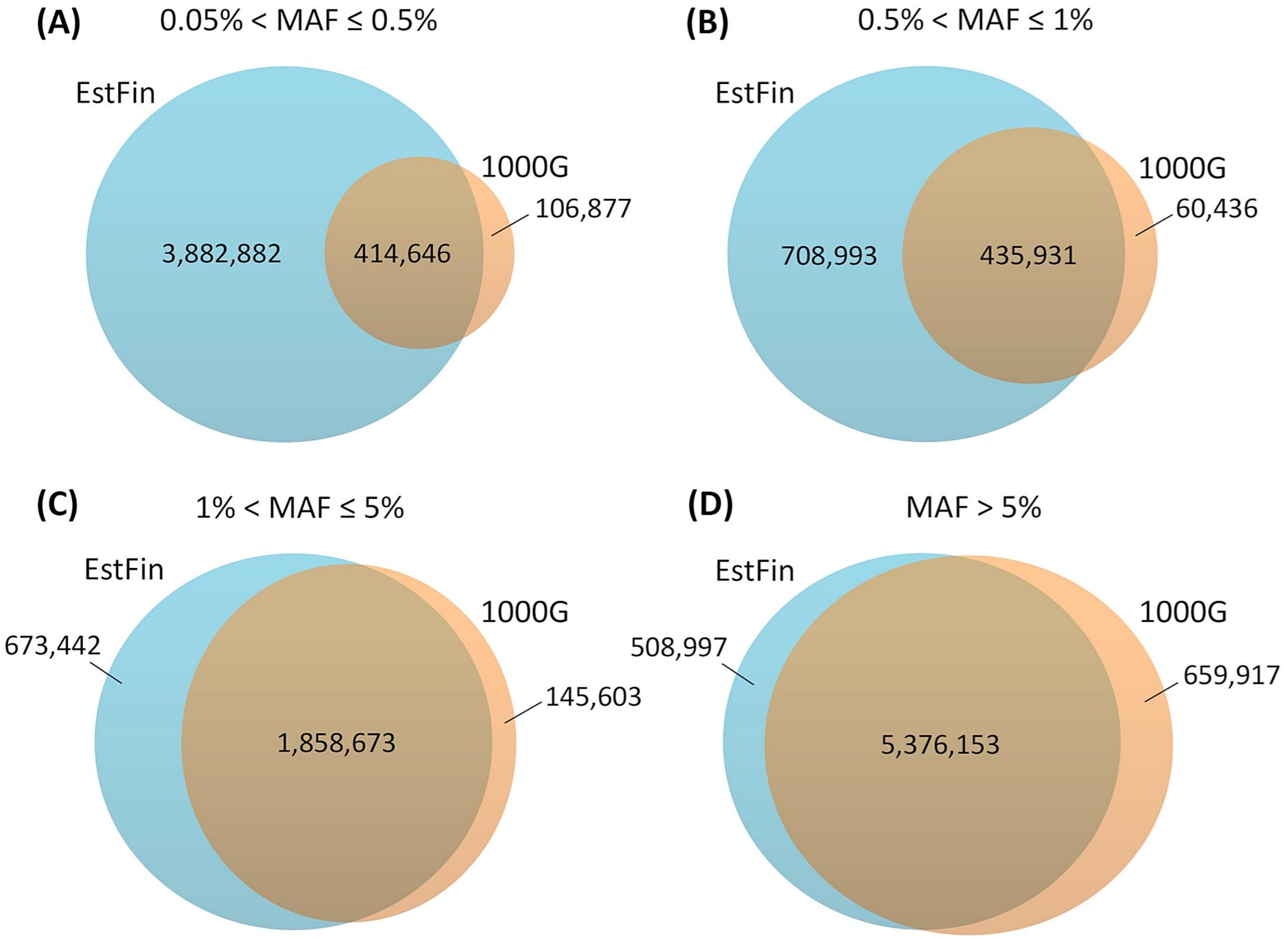

### S2 Figure

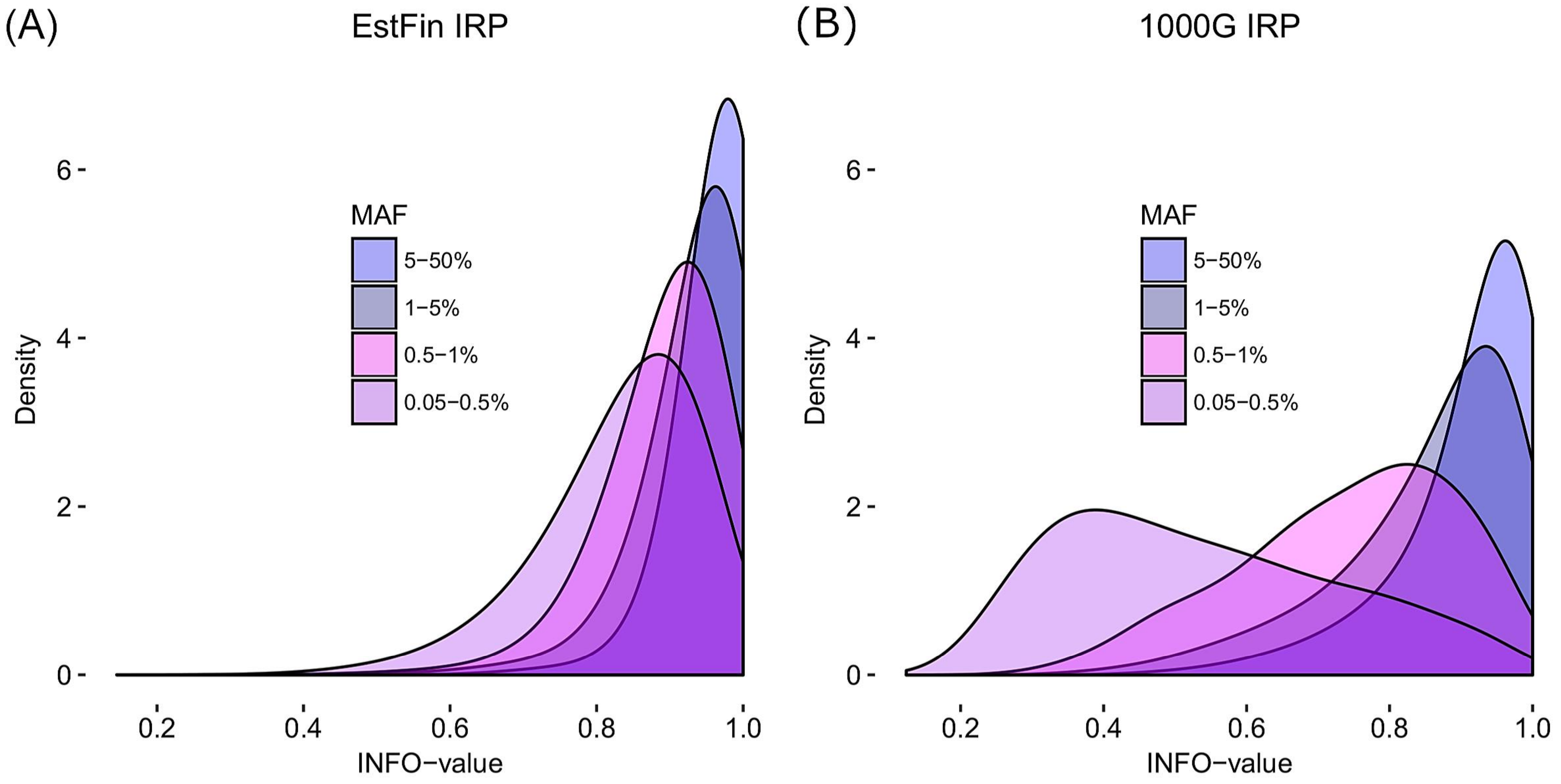

### S3 Figure

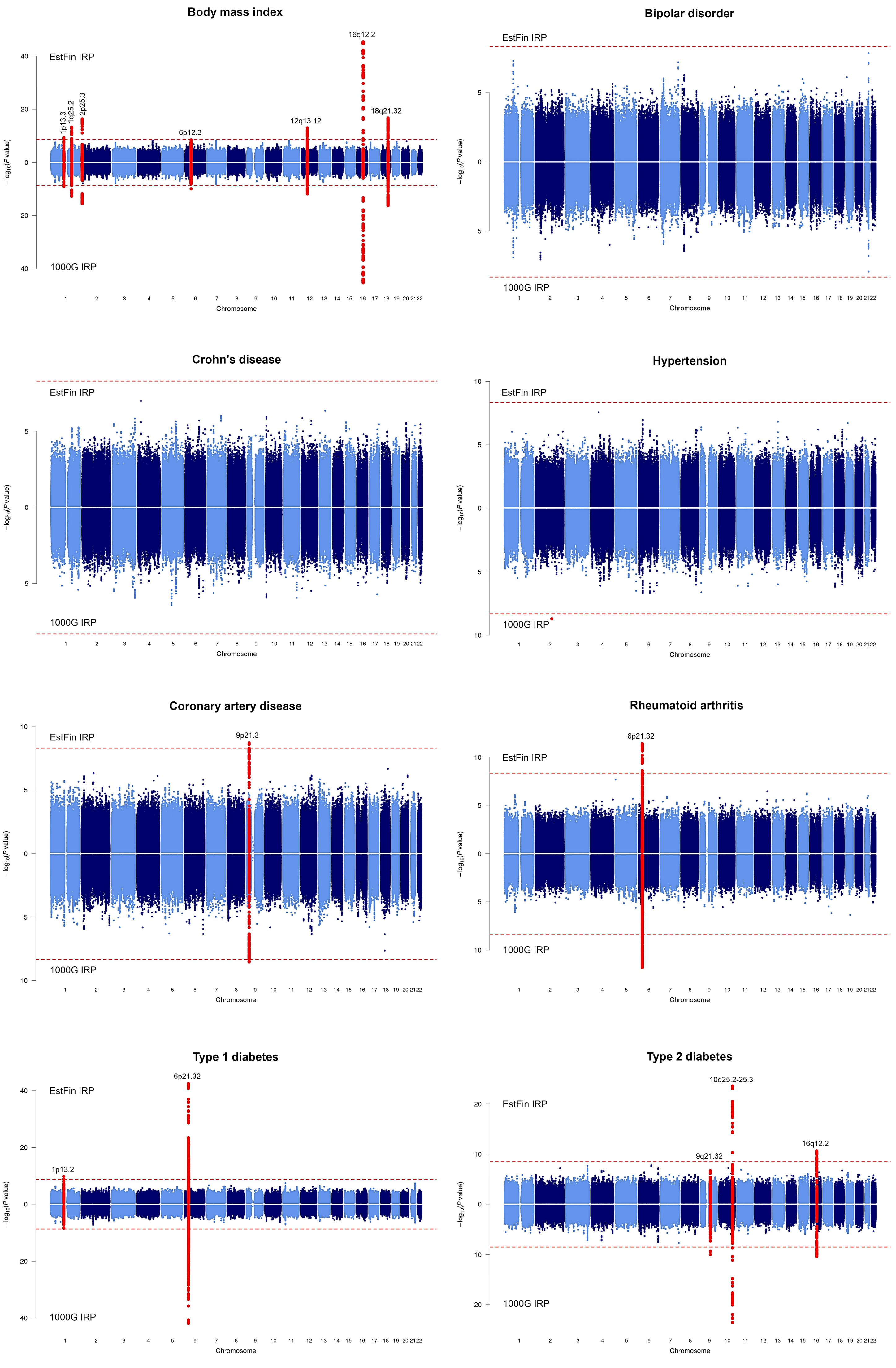

### S4 Figure

#### Body mass index at 1p13.3

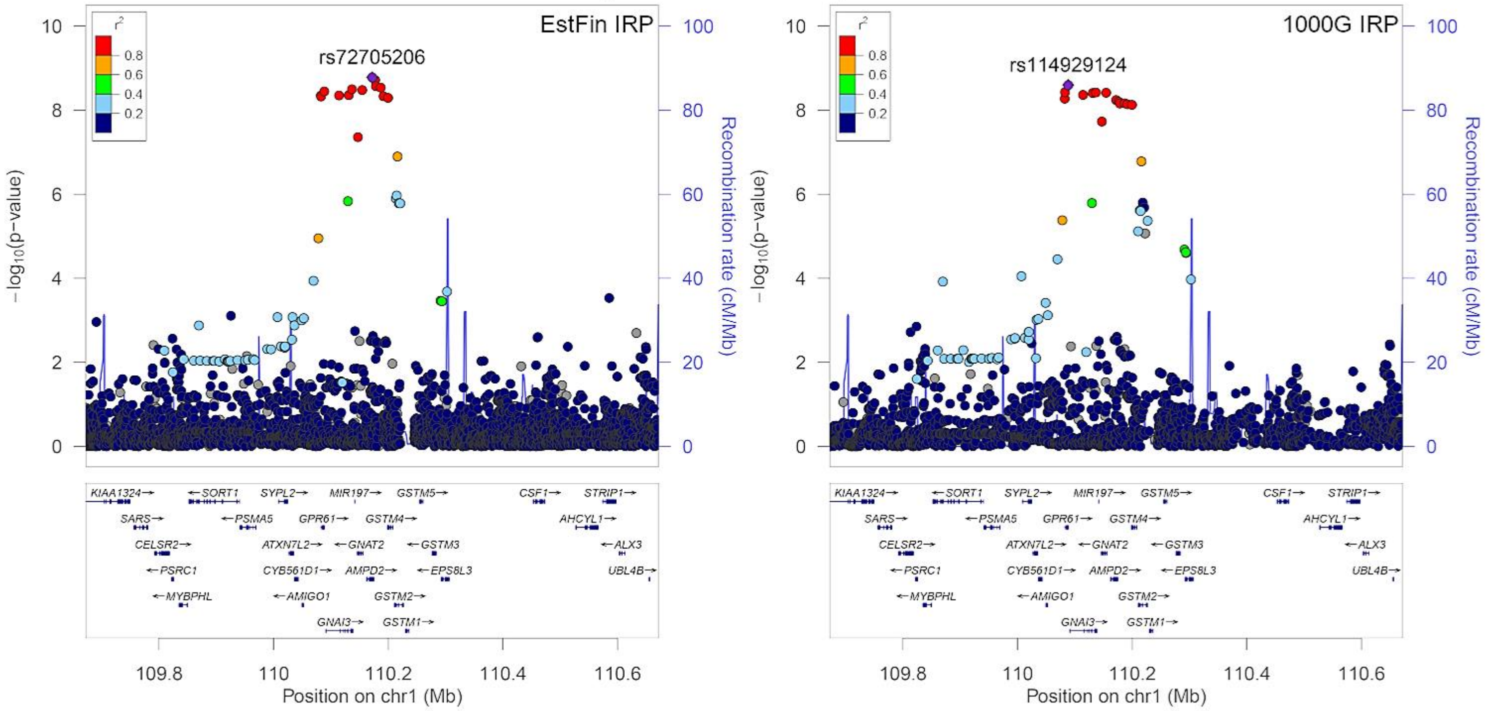


#### Body mass index at 1q25.2

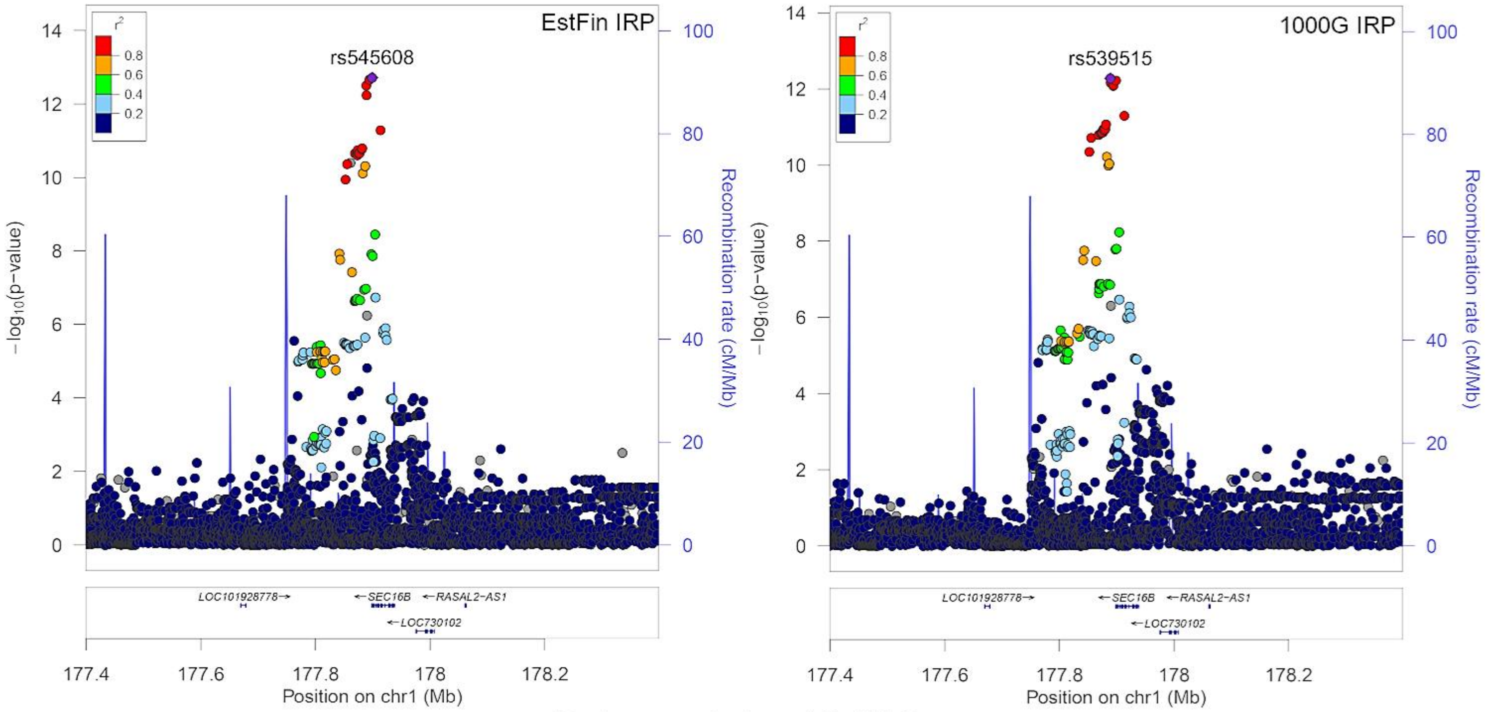


#### Body mass index at 2p25.3

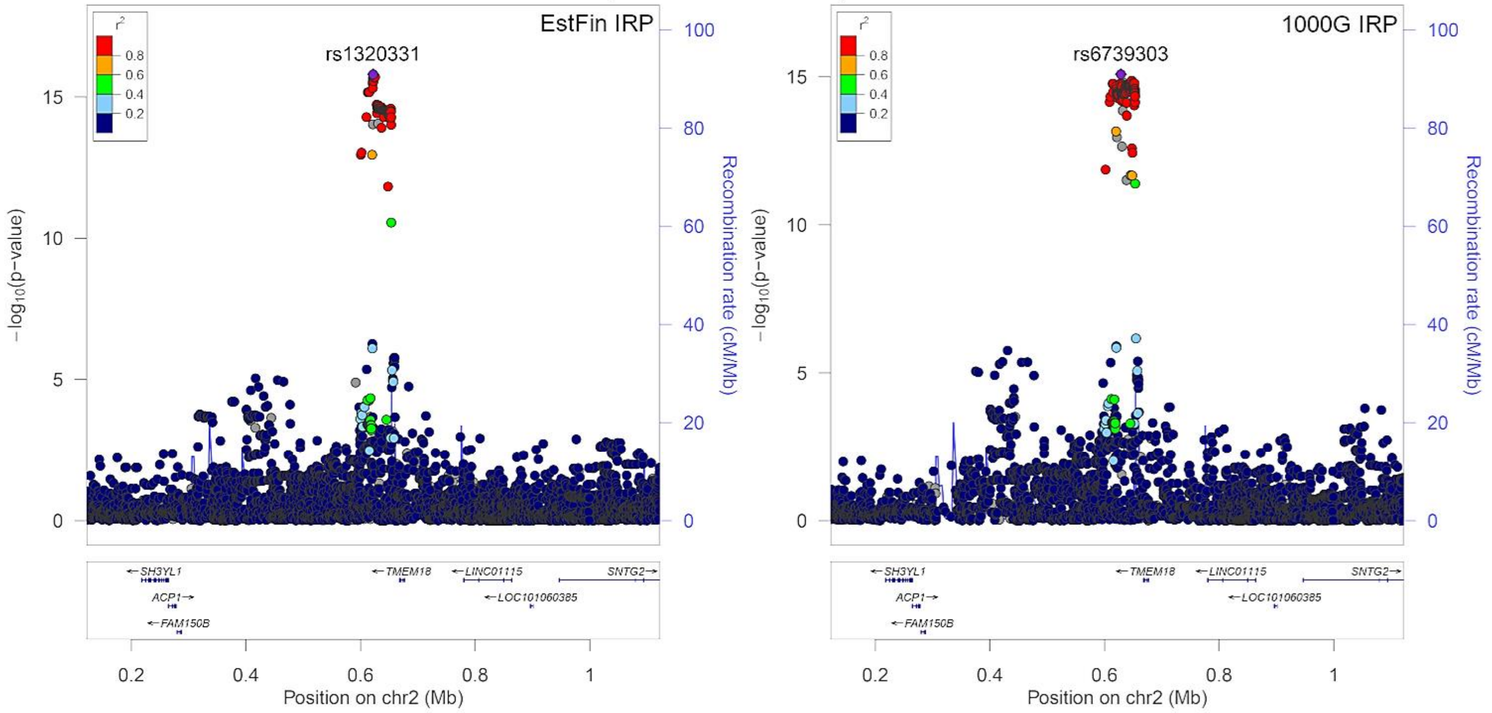

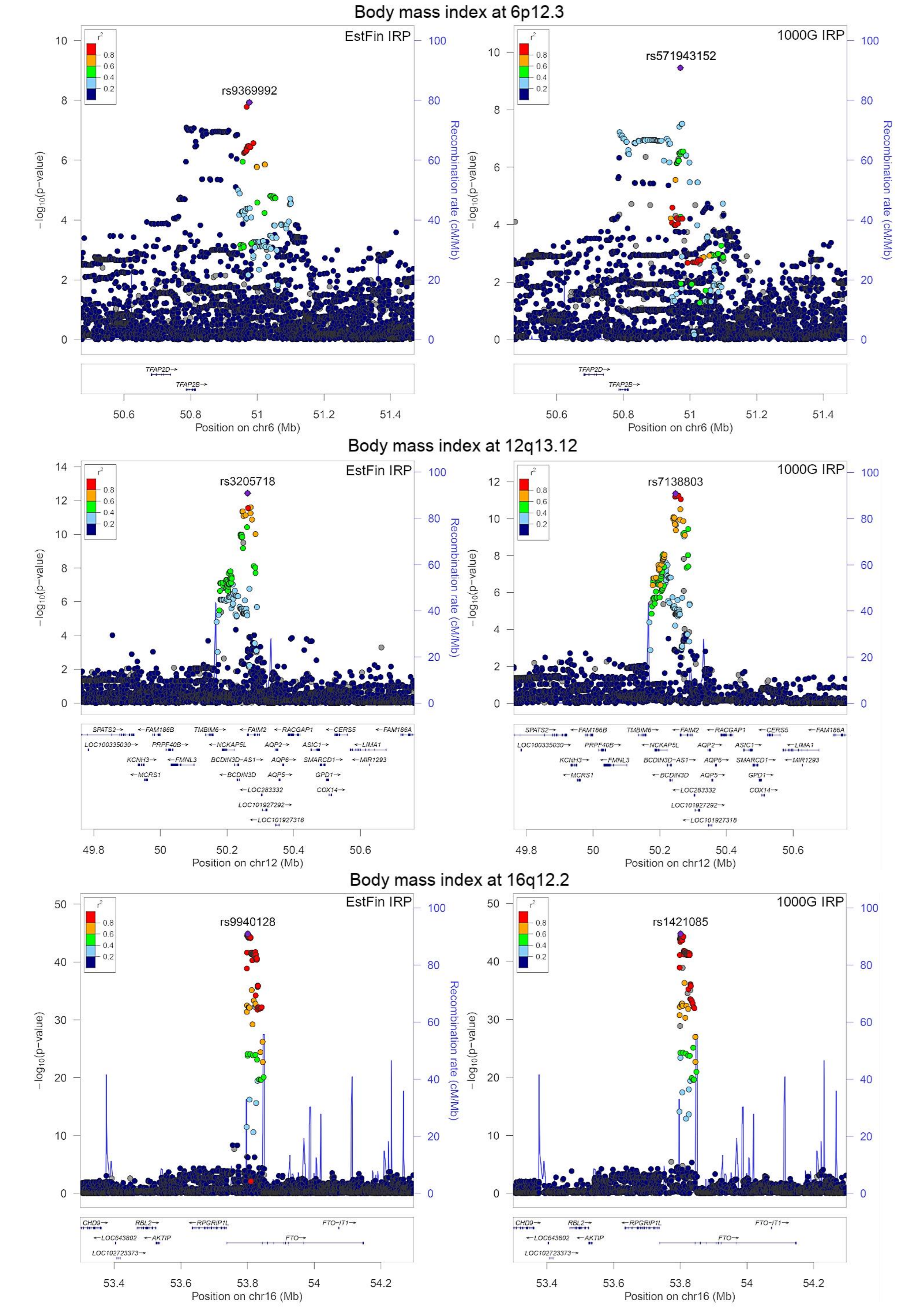

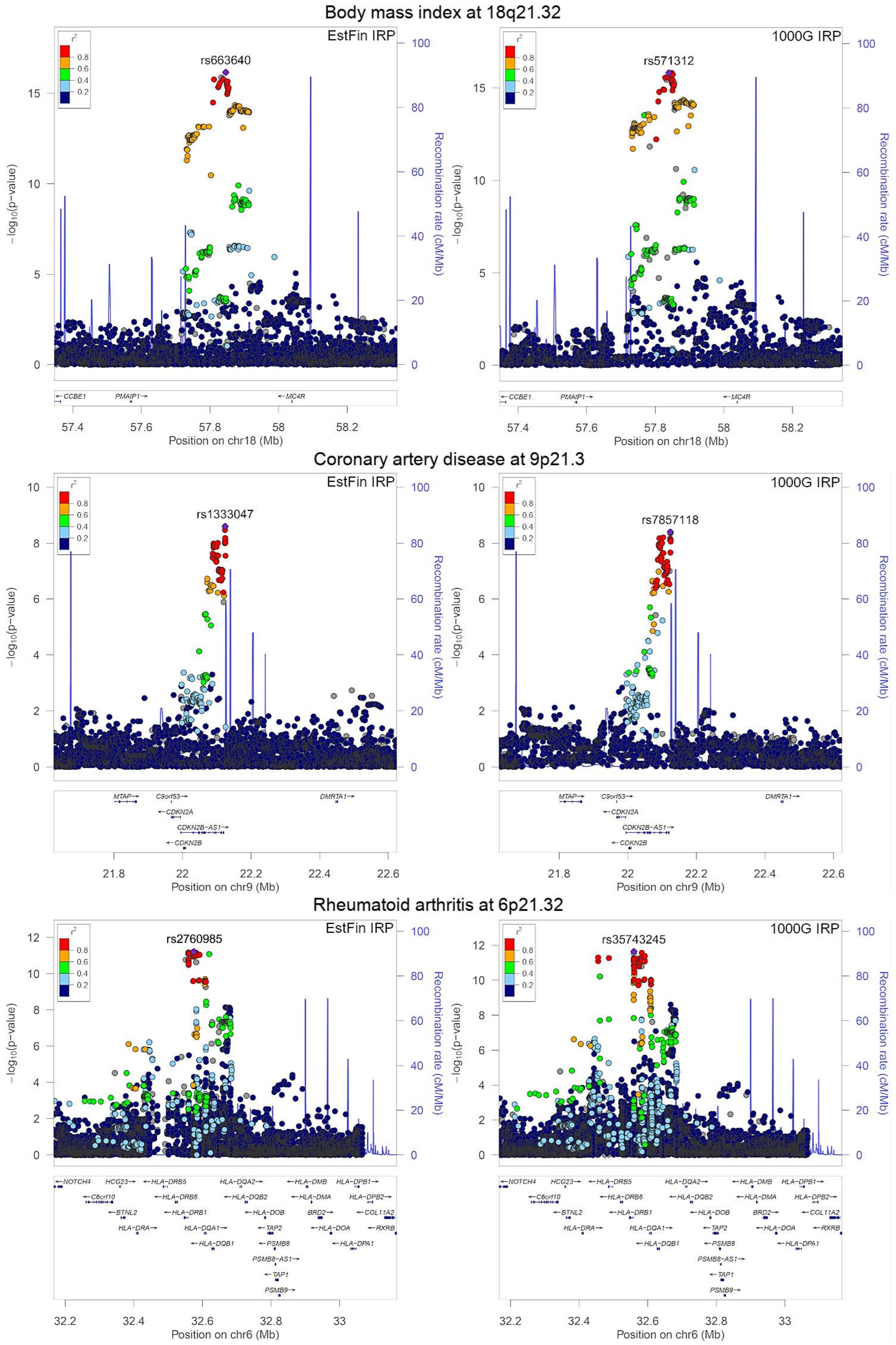

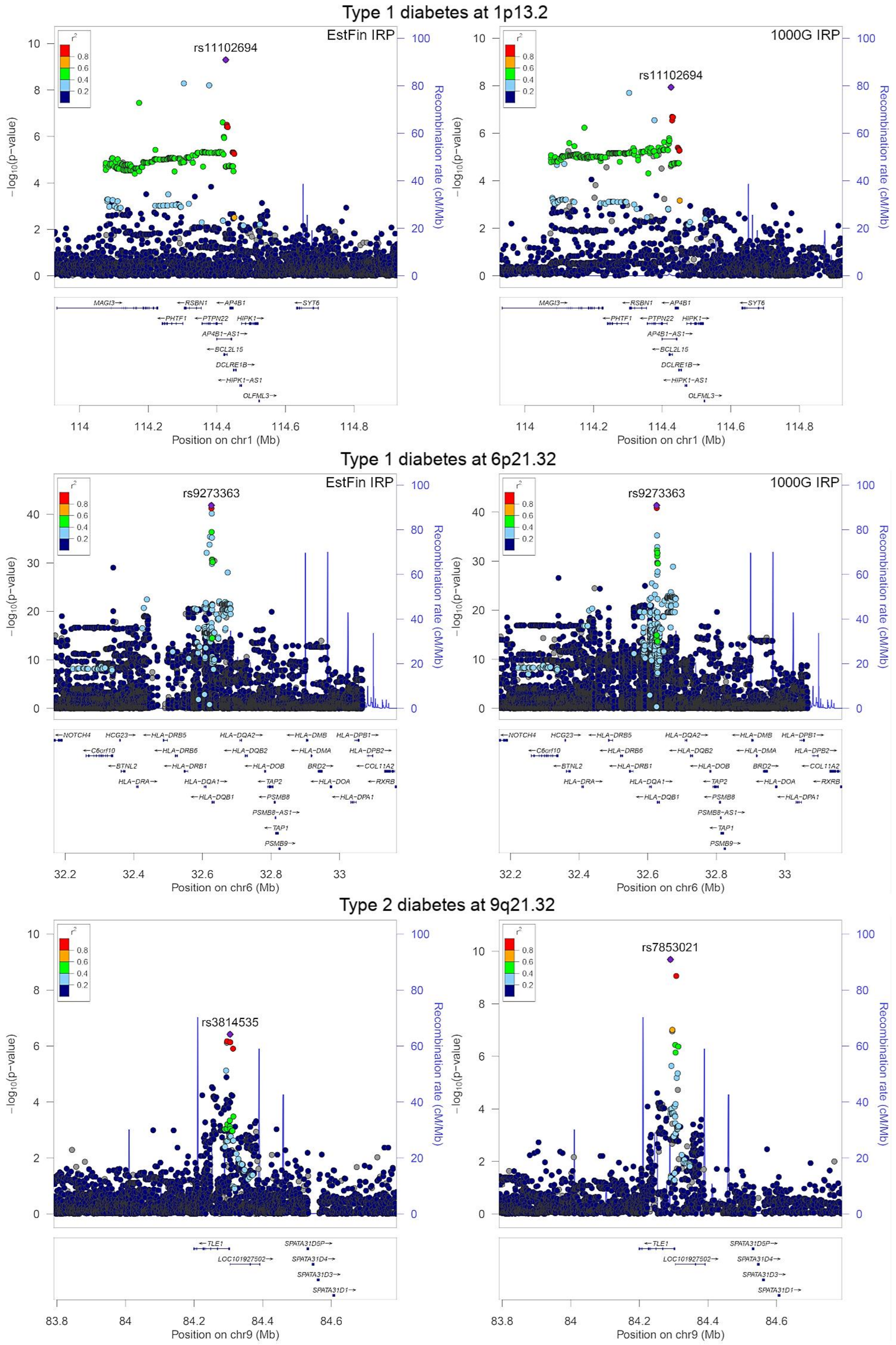

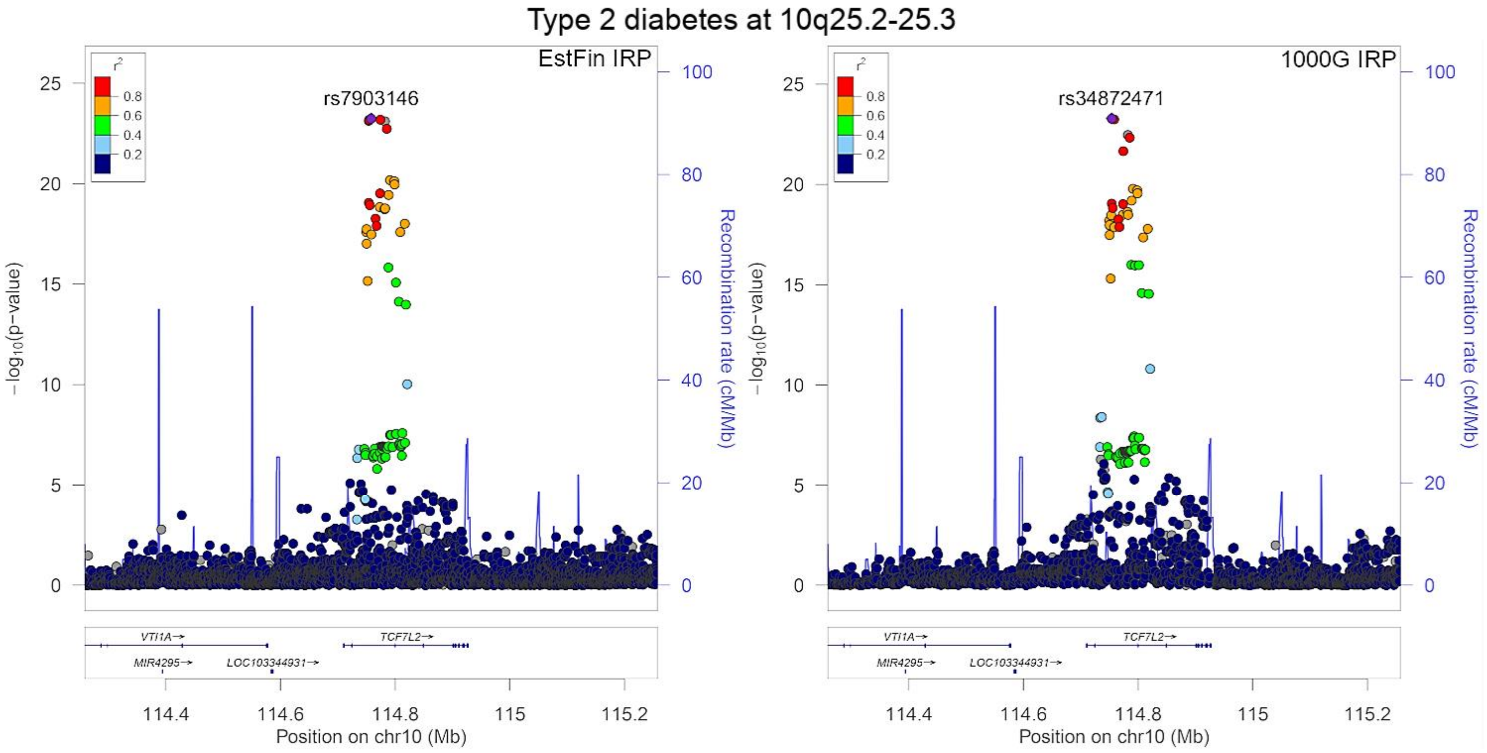

### S5 Figure

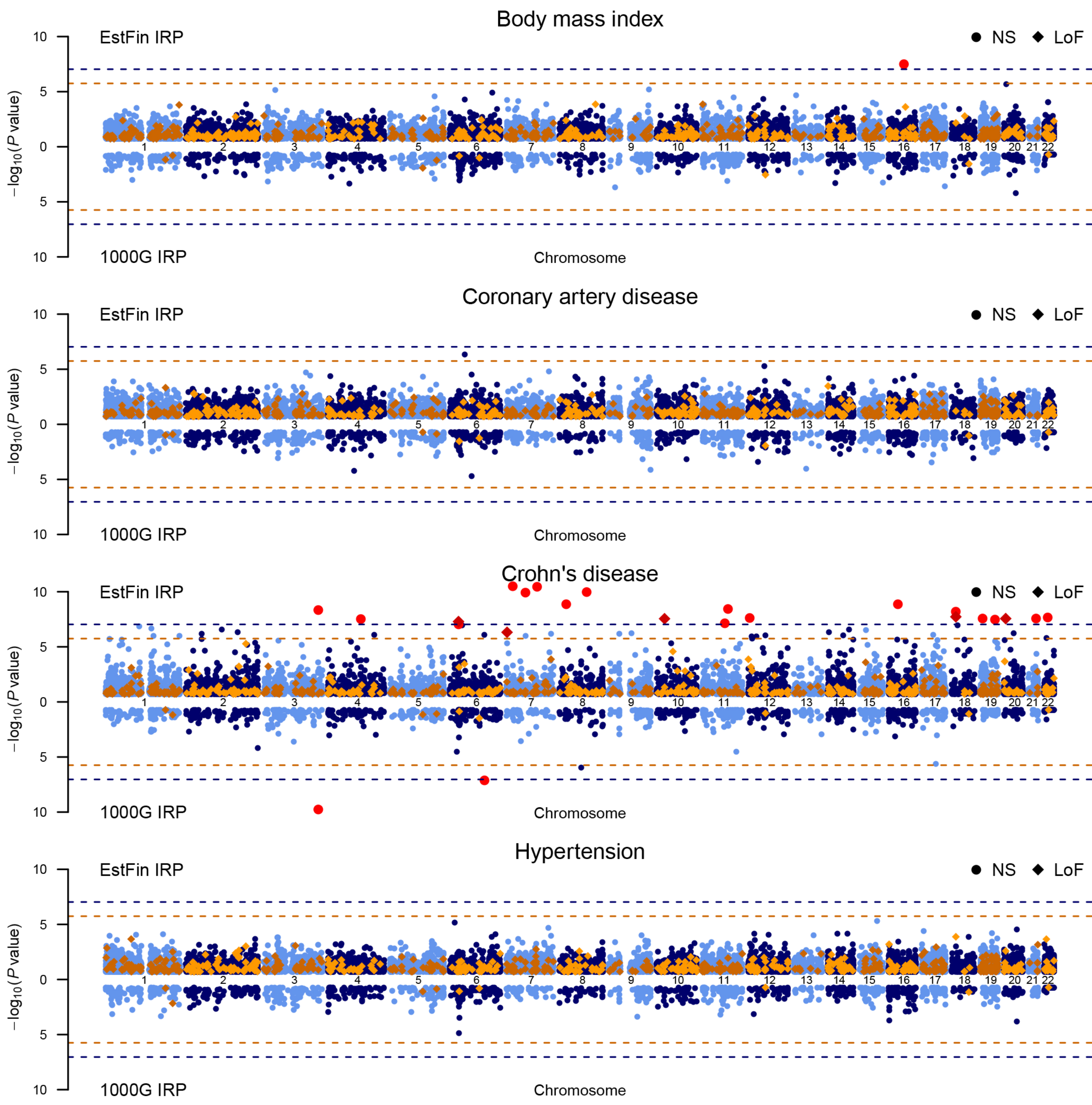

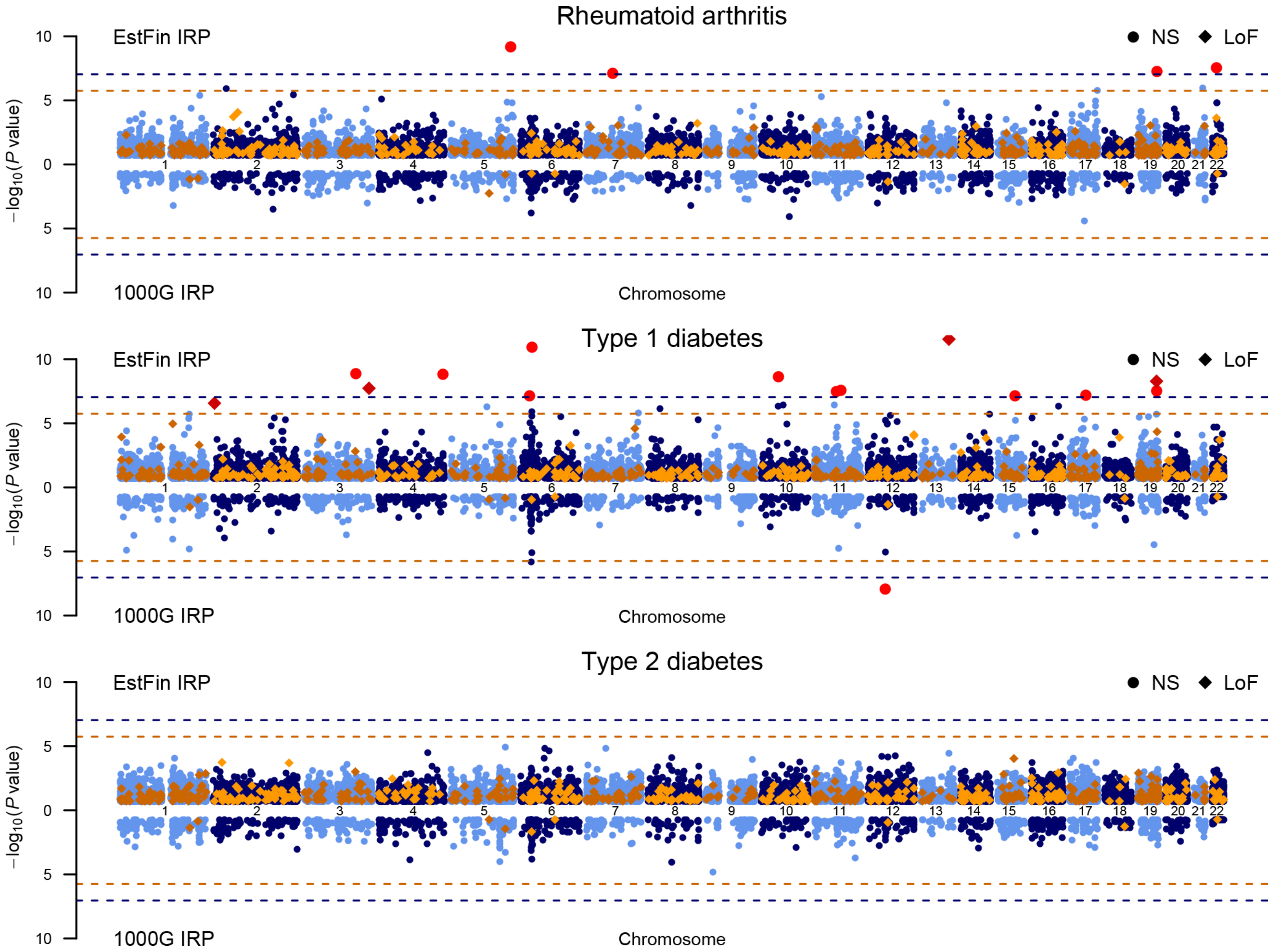

### S6 Figure

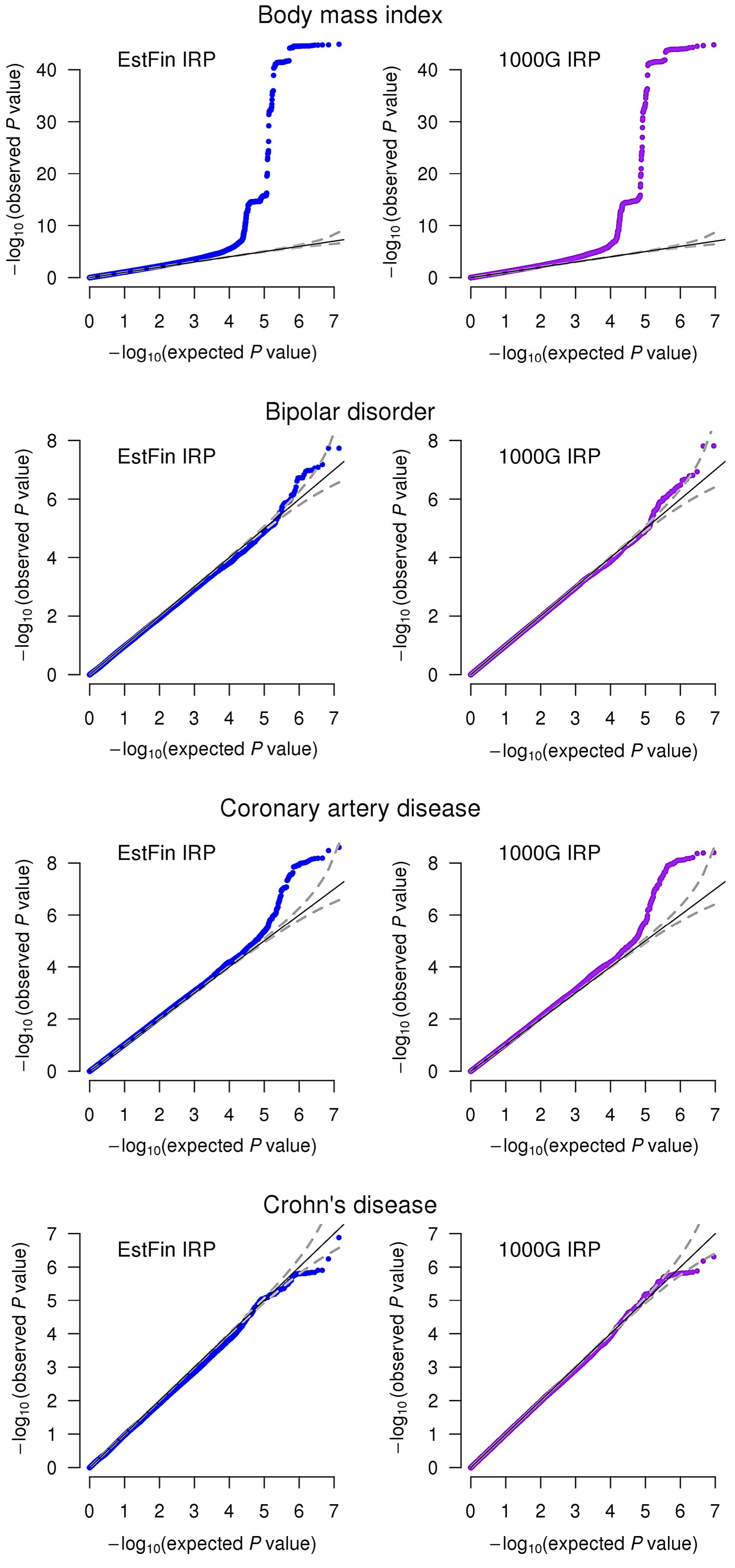

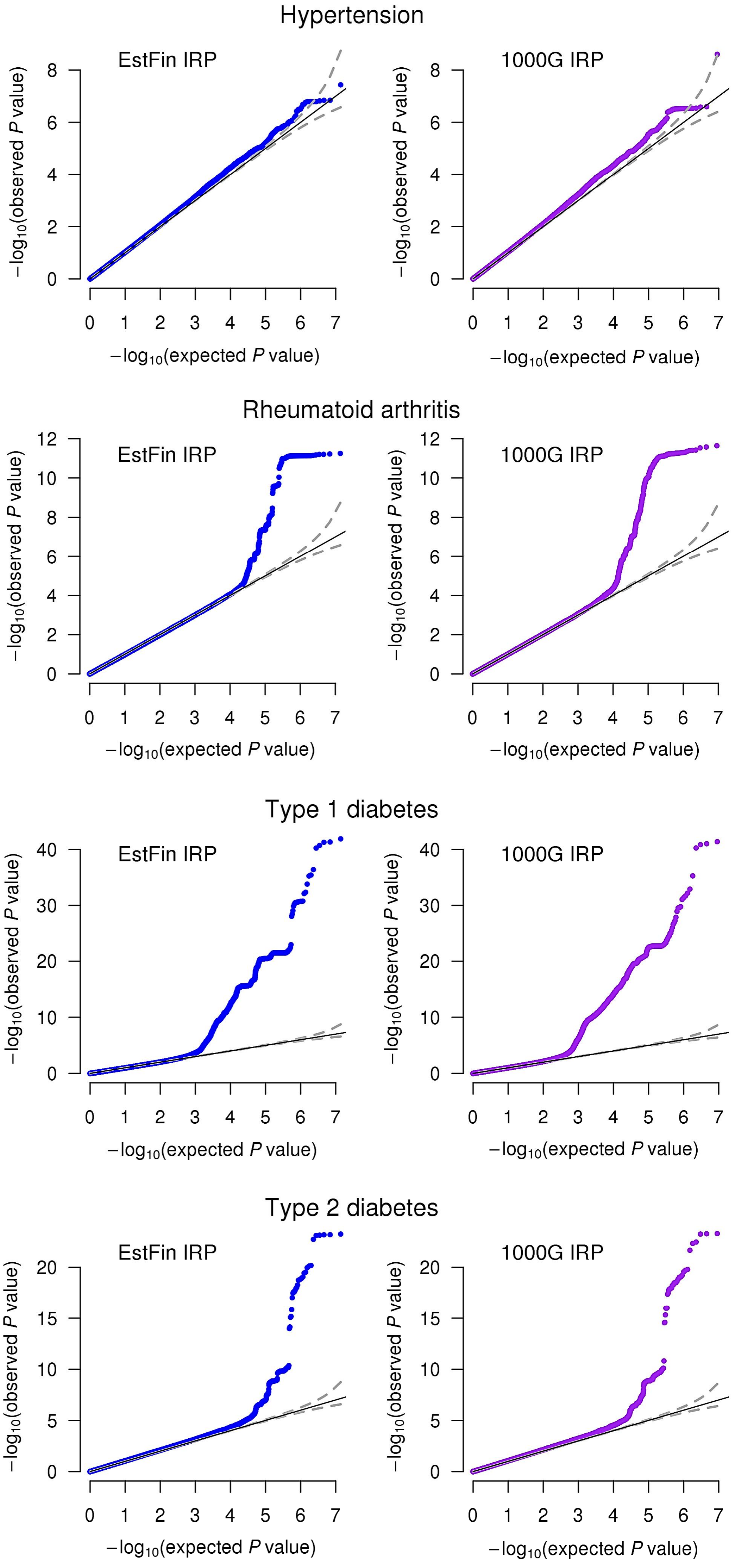
