## Supplementary material for "Advantages of genotype imputation with ethnically matched reference panel for rare variant association analyses": S1 Appendix

### Genotype imputation

In the genome-wide association analyses, we studied 13,859,717 and 9,058,236 confidently imputed (CI) variants (INFO-value > 0.8) with MAF > 0.05% based on the EstFin and the 1000G IRPs in 36,716 unrelated individuals, respectively. Both imputed datasets shared 8.09 M CI variants, but 5.77 M were specific to the EstFin-based imputation and 0.97 M to the 1000G-based data (S1 Table and S1 Fig). We observed many more (4.3 M vs. 0.5 M) rare (0.05 < MAF ≤ 0.5%) and moderately more (3.7 M vs. 2.5 M) low-frequency (0.5 < MAF ≤ 5%) CI variants in the data imputed with the EstFin IRP compared to the 1000G-based data. Large number (5.38 M) of common CI variants (MAF > 5%) overlapped in both imputed datasets.

As shown before [1], both IRPs provided lower imputation quality for lower frequency variants (S2 Fig). For the EstFin IRP, the improvement of imputation confidence at each MAF category is almost homogenous. Controversially, diverse 1000G panel has a tendency to employ incorrect allele frequency distribution – only ~15% of rare (0.05 < MAF ≤ 0.5%) and ~69% of low-frequency (0.5 < MAF ≤ 5%) variants are confidently imputed. Majority (~99% with the EstFin IRP and ~90% with the 1000G IRP) of common variants (MAF > 5%) are confidently imputed in both datasets.

### Disease associated genes

#### Bipolar disorder: *VGLL4* (NS variants)

Variants in *VGLL4* have shown associations with anorexia nervosa [2,3], schizophrenia [4], and with impulsivity facets of neuroticism [5]. An overlap in genetic predisposition for neuroticism, bipolar disorder, major depressive disorder and schizophrenia has been delineated [6]. *VGLL4* encodes vestigial-like 4, a protein that regulates gene expression in myocytes and acts as a tumour suppressor in gastric cancer through counteracting oncogenes. Its function in the brain has not yet been described. In Wnt/β-catenin signalling pathway, *VGLL4* negatively regulates Wnt/β-catenin signalling pathway via inhibiting β-catenin and TCF (T-cell factor). *VGLL4* can also suppress epithelial-mesenchymal transition (EMT) and contribute to apoptosis signalling pathway.

### Crohn's disease: *HLA-G* (NS/LoF variants)

We replicated known associations of *HLA-G* variants with Crohn's disease [7]. *HLA-G* is involved in Class I MHC mediated antigen processing and presentation and cytokine signalling in immune system. The loss of *HLA-G* mediated control of the immune responses may lead to the onset of autoimmune/inflammatory diseases, caused by an uncontrolled activation of the immune effector cells and important role of this molecule in the modulation of the immune system has been proposed [8].

### Crohn's disease: *FERMT1* (LoF variants)

Association of *FERMT1* loss-of-function variants with Crohn's disease gives support to findings from the whole exome sequencing study in pediatric IBD patients postulating that in a subset of IBD, heterozygous mutations in genes presumed to manifest IBD following the autosomal recessive inheritance pattern may contribute to clinical presentation [9]. Generally, loss-of-function mutations in the *FERMT1* gene encoding the focal adhesion protein, fermitin family homolog-1 are resulting in Kindler syndrome, an autosomal recessive disorder characterized by skin atrophy and blistering. Our variants are not listed in ClinVar [10] and could probably be population-specific.

### Rheumatoid arthritis: *HSPB1* (NS variants)

*HSPB1* encodes a molecular chaperone Hsp27, a member of the small heat shock protein (HSP20) family of proteins (sHsp) which are constitutively expressed in response to a wide variety of unfavourable physiological and environmental conditions and sharing functions with protection against toxicity mediated by aberrantly folded proteins or oxidative-inflammation conditions [11,12]. Extracellular sHsps are implicated in cell-cell communication, activation of immune cells, and promoting anti-inflammatory and anti-platelet responses [13].

The spontaneous production and release of TNF, IL-1β, IFNγ and IL-8 is a key feature of the synovial tissue in RA [14]. HSP27 expression is required for IL-1-induced expression of the pro-inflammatory mediators cyclooxygenase-2, IL-6, and IL-8. Likewise, it is required for the activation of both IL-1 and TNF-induced signalling pathways for which the most upstream common signalling protein is transforming growth factor-beta-activated kinase-1 (TAK1), and for IL-1 TAK1-mediated downstream signalling by p38 MAPK, JNK, and their activator [15].

Toll-like receptors (TLRs) are innate immune receptors that respond to both exogenous and endogenous stimuli and are suggested to contribute to the perpetuation of chronic inflammation associated with rheumatoid arthritis pathogenesis, particularly the endosomal TLRs 3, 7, 8, and 9 [16]. Inhibition of TLR2 signalling has been proposed to block the perpetuation of an inflammation cycle driven by endogenous ligands being recognized by TLR2 [14]. A paralog of *HSPB1* – *HSPB8* – is recognised as an endogenous TLR4 ligand being abundantly expressed in RA synovial tissue and a role in the inflammatory process in autoimmune diseases such as RA was suggested [17].

#### Type 1 diabetes: *LPP* (LoF variants)

Variants in *LPP* have shown associations with diverse autoimmune diseases (pleiotropic effect) such as celiac disease [18], juvenile idiopathic arthritis [19], multiple sclerosis [20], systemic lupus erythematosus [21], and on the other hand, chronic lymphocytic leukemia [22] and type 2 diabetes [23]. We report a joint association of rare loss-of-function variants in *LPP* with T1D contradictory to findings from where *LPP* as celiac disease susceptibility gene showed no evidence of association with type 1 diabetes [18]. However, *LPP* mRNA and protein are expressed in multiple tissues, including islets of Langerhans and pancreas and *LPP* gene is relatively intolerant of loss-of-function variation (ExAC pLI = 0.58) [24].
